# Genealogical analysis of replicate flower colour hybrid zones in *Antirrhinum*

**DOI:** 10.1101/2025.04.27.649910

**Authors:** Arka Pal, Daria Shipilina, Alan Le Moan, Adrian J. McNairn, Jennifer K. Grenier, Marek Kucka, Graham Coop, Yingguang Frank Chan, Nicholas H. Barton, David L. Field, Sean Stankowski

**Affiliations:** Institute of Science and Technology Austria, 3400 Klosterneuburg, Austria; UMR 7144 AD2M, CNRS-Sorbonne Université, Station Biologique de Roscoff, Roscoff, France; Genomics Innvoation Hub and TREx Facility, Cornell University, Ithaca, NY, USA; Friedrich Miescher Laboratory of the Max Planck Society, 72076 Tübingen, Germany; Department of Translational Genomics, University of Cologne, 50931 Cologne, Germany; Department of Evolution & Ecology and Center for Population Biology, University of California – Davis, CA, USA; Groningen Institute for Evolutionary Life Sciences (GELIFES), University of Groningen, 9747 AG Groningen, the Netherlands; Applied BioSciences, Macquarie University, Sydney, New South Wales, Australia; Department of Ecology and Evolution, The University of Sussex, Brighton, United Kingdom

**Keywords:** Ancestral Recombination Graphs, genealogy, barriers to gene flow, flower colour, snapdragon, hybrid zones

## Abstract

A major goal of speciation research is identifying loci that underpin barriers to gene flow. Population genomics takes a ‘bottom-up’ approach, scanning the genome for molecular signatures of processes that drive or maintain divergence. However, interpreting the ‘genomic landscape’ of speciation is complicated, because genome scans conflate multiple processes, most of which are not informative about gene flow. However, studying replicated population contrasts, including multiple incidences of secondary contact, can strengthen inferences. In this paper, we use linked-read sequencing (haplotagging), *F*ST scans, and genealogical methods to characterise the genomic landscape associated with replicate hybrid zone formation. We studied two flower colour varieties of the common snapdragon, *Antirrhinum majus* subspecies *majus*, that form secondary hybrid zones in multiple independent valleys in the Pyrenees. Consistent with past work, we found very low differentiation at one well-studied zone (Planoles). However, at a second zone (Avallenet), we found stronger differentiation and greater heterogeneity, which we argue is due to differences in the amount of introgression following secondary contact. Topology weighting of genealogical trees identified loci where haplotype diversity was associated with the two snapdragon varieties. Two of the strongest associations were at previously identified flower colour loci: *Flavia*, that affects yellow pigmentation, and *Rosea/Eluta*, two linked loci that affect magenta pigmentation. Preliminary analysis of coalescence times provides additional evidence for selective sweeps at these loci and barriers to gene flow. Our study highlights the impact of demographic history on the differentiation landscape, emphasizing the need to distinguish between historical divergence and recent introgression.

## 1 | Introduction

A major goal of speciation research is to identify loci underlying barriers to gene flow. Population genomic studies usually take a ‘bottom-up’ approach by scanning the genome for patterns of within- and between-population variation that indicate selection driving or maintaining divergence (Ravinet et al., 2017; Wolf & Ellegren, 2017). For example, during speciation with gene flow, genomic regions associated with local adaptation or genetic incompatibilities are expected to show elevated genetic differentiation (usually measured by *F*ST), with the rest of the genome homogenised through genetic exchange (Feder et al., 2012; Wu, 2001). Indeed, numerous studies of the ‘genomic landscape’ have found highly heterogenous patterns of genetic differentiation and, in some cases, have shown that regions with high *F*ST house genes underpinning adaptive traits that also act as reproductive barriers (Hooper et al., 2024; Martin et al., 2013; Poelstra et al., 2014; Todesco et al., 2020). However, we now know that interpreting the differentiation landscape is more challenging than some researchers once hoped (Ravinet et al., 2017; Wolf & Ellegren, 2017).

The main challenge is that genome scans can conflate multiple processes, some of which are not directly relevant to current heterogenous gene flow (Ravinet et al., 2017). Consider a simple model of secondary contact, where divergence builds up over a long period (which may involve intermittent isolation), and erodes following recent contact. Genome-wide divergence builds up relatively slowly, due to both drift and selection. Divergence will inevitably be heterogenous along the genome both by chance and due to intrinsic properties of the genome, such as the local density of functional elements and local recombination rate (Burri, 2017). After contact, introgression will erode divergence where the populations meet, potentially revealing the location of barrier loci (Duranton et al., 2018). While relatively fast compared to build-up of divergence, this erosion takes some time, and will be delayed if interbreeding is geographically localised (Barton & Gale, 1993). Thus, genome scans reflect both initial divergence and post-contact introgression, and these may be hard to disentangle.

Inclusion of replicate hybrid zones aids the interpretation of genome scans, allowing comparison of divergence across multiple contacts (Nadeau et al., 2014; Rancilhac et al., 2024; Vijay et al., 2016; Wilding et al., 2001). Overall divergence may reflect differences in the timing of contact or rates of gene flow. Nevertheless, parallel contacts should ultimately lead to similar differentiation landscapes if large-effect outlier loci reflect barriers to gene flow that have resisted introgression in each location. In contrast, outliers found in a single zone might reflect local demographic processes (e.g., bottlenecks), evolutionary noise, sampling effects, or population specific barriers (Westram et al., 2021). Several studies have used this logic to identify loci that underpin local adaptation and speciation. The most compelling studies combine traditional site-based genome scans with tree-based methods, which make it possible to analyse more than two populations within a single framework that acknowledges their recent shared history (Poelstra et al., 2014; Rancilhac et al., 2024).

In this paper, we study genome-wide variation associated with replicate hybrid zones in the common snapdragon, *Antirrhinum majus*, a classic model for understanding phenotypic variation both in the laboratory and in nature (Hudson et al., 2008). We focus on two varieties of *A. majus* subspecies *majus—A.m.m* var. *pseudomajus* and *A.m.m* var. *striatum* (hereafter, var. *pseudomajus* and var. *striatum* for brevity)*—*that are native to France and Spain (Whibley et al., 2006). These varieties have largely non-overlapping geographic distributions, occupy similar habitats and are pollinated by the same bee species (Tavares et al., 2018). The major difference between them is their contrasting flower colour: var. *pseudomajus* has magenta flowers with a small patch of yellow pigment on the face of the flower below the bee entry point, while var. *striatum* has yellow with restricted veins of magenta coloration above the bee entry point (Fig. 1A). These differences in colour, which are thought to be alternative adaptations to attract the same bee pollinators, are caused by a small number of loci that control the production of two flavonoid pigments in floral tissue, anthocyanin (magenta) and aurone (yellow). *Rosea*, *Eluta* (Tavares et al., 2018), *and Rubia* (Field et al., 2025) affect anthocyanin production, while, *Sulfurea* (Bradley et al., 2017), *Flavia* (Bradley et al., 2025), *Cremosa* and *Aurina* (Richardson et al., 2025) affect aurone production.

**Figure 1.**
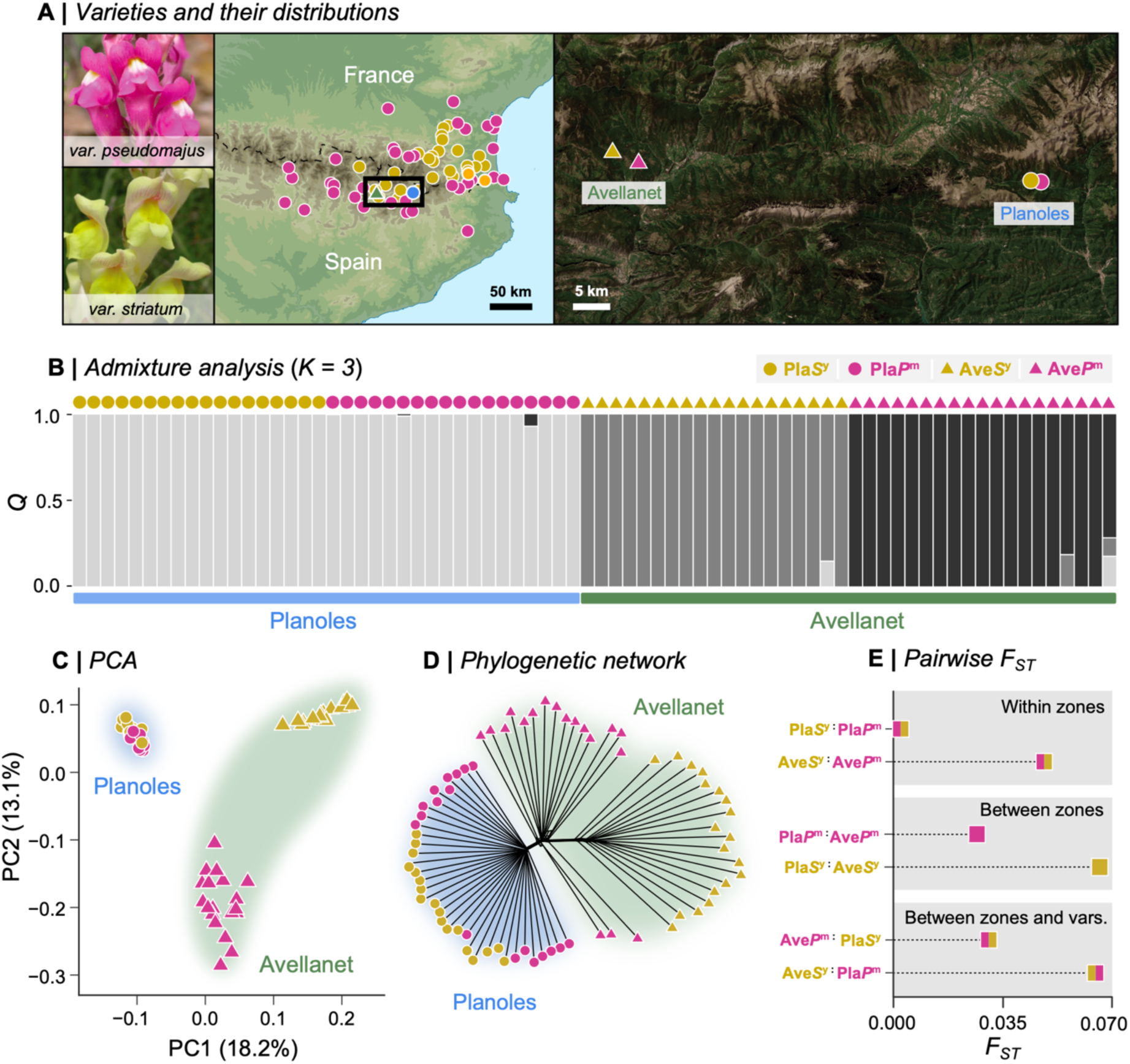
Evolutionary relationship between *A. majus* subspecies *majus* populations from two hybrid zones. **(A)** Geographic distributions of magenta flowered *A.m.m.* var. *pseudomajus* and yellow-flowered *A.m.m.* var. *striatum*. The map is based on sample locations in Whibley et al. (2006) and does not show the full distribution of either variant. Circles are coloured according to the population. Samples were collected from two hybrids zones: Avellanet (19 magenta & 19 yellow samples) and Planoles (18 magenta & 18 yellow samples). Points on the map represent the average location of each population. **(B)** Genetic structure, shown as admixture coefficient (*Q*) for *K*=3 clusters inferred by *Admixture* from 1.7 million LD-thinned SNPs. Each vertical bar is one individual. **(C)** The first two principal components of the same dataset. **(D)** Phylogenetic network (*neightbourNet*) on the same dataset. **(E)** Estimates of per-site Weir & Cockerham’s *F*_ST_, averaged over all 11.5 million SNPs. *F*_ST_ was calculated between varieties from the same hybrid zone (Pla*P*^m^ vs Pla*S*^y^, Ave*P*^m^ vs Ave*S*^y^), and between the hybrid zones (Pla*P*^m^ vs Ave*P*^m^, Pla*S*^y^ vs Ave*S*^y^, Pla*S*^y^ vs Ave*P*^m^, Pla*P*^m^ vs Ave*S*^y^). Pla: Planoles, Ave: Avellanet, *P*^m^: magenta-coloured var. *pseudomajus, S*^y^: yellow-coloured var. *striatum*.

During the last ice age, var. *pseudomajus* and var. *striatum* are thought to have been restricted to areas of low elevation, but subsequently expanded into the Spanish Pyrenees (Vargas et al., 2004; Whibley et al., 2006). As a result, at least three separate hybrid zones have formed in separate valleys below the altitudinal limit of *A. majus* (Fig. 1A). In one such zone near the town of Planoles, a transition from yellow to magenta flowers occurs over a few kilometres (Whibley et al., 2006). Scans of genome-wide sequence variation have revealed strong allele frequency differentiation and sharp geographic clines around previously identified colour loci (Field et al., 2025; Tavares et al., 2018). In contrast, most of the surrounding genome shows low genetic differentiation, probably owing to the homogenising effects of dispersal and recombination (Ringbauer et al., 2018; Tavares et al., 2018).

Here, we expand our analysis to include individuals from the Planoles hybrid zone and a second zone near the town of Avellanet, located over 50 km to the west (Fig. 1A). By comparing their respective genomic landscapes and jointly analysing two independent hybrid zones, we hoped to disentangle ancestral divergence from the effects of recent introgression. We were especially interested in whether known flower colour loci act similarly in both localities and stand out from their genomic background. To address this, we used both traditional *F*ST scans, and genealogical methods for studying the genome-wide distributions of tree topologies and coalescence times across the genome. As a secondary aim, we use this study as an opportunity to compare different methods for inferring genealogical trees along the genome. Several approaches are now available for inferring genealogies from phased SNP datasets (Nielsen et al., 2024). However, these methods have not yet been widely used to study adaptation and speciation. It is also unclear how they perform when applied to real datasets, and there has been limited discussion about when more sophisticated methods might be warranted over simpler ones. We hope that this study helps other researchers decide which method might be most appropriate for their data and specific goals.

## 2 | Results and Discussion

### 2.1 | Genome-wide analysis reveals different histories of post-contact gene flow across the two hybrid zones

We sampled 18 individuals of magenta-coloured var. *pseudomajus* and 18 yellow-coloured var. *striatum* from Planoles (hereafter, Pla*P*^m^ and Pla*S*^y^), as well as 19 of each variety from Avellanet (hereafter, Ave*P*^m^ and Ave*S*^y^) (Table S1). We sequenced them using haplotagging, a method of linked-read sequencing (Meier et al., 2021). An advantage of haplotagging over standard short-read sequencing is the ability to track source haplotypes by means of molecular barcoding. After mapping sequence reads to the *A. majus* reference genome v. 3.5 (Li et al., 2019), we followed the variant calling and imputation pipeline outlined in Meier et al. (2021) that leverages linked-read information to identify 11,533,030 bi-allelic SNPs (22 SNPs/kbp) across all of the samples. We then phased the SNPs using *SHAPEIT5* (Hofmeister et al., 2023) and used information from a closely related outgroup (*A. molle*; Durán-Castillo et al., 2022) to polarise variants as ancestral or derived.

Based on previous work that showed low genome-wide differentiation between the varieties in Planoles (Tavares et al., 2018), we expected to see similarly low differentiation at the previously unstudied hybrid zone at Avellanet. To test this hypothesis, we generated a LD-thinned dataset containing 1.71 million SNPs and performed *Admixture* (Fig 1B; S1), principal component (Fig. 1C) and phylogenetic (Fig. 1D, S2) analysis to characterise genetic structure. In contrast to our expectations, we found different patterns of genetic structure at each hybrid zone. Specifically, Pla*P*^m^ and Pla*S*^y^ always formed a single group, rather than clustering by flower colour. In contrast, the Ave*P*^m^ and Ave*S*^y^ always formed two distinct groups. This result was also supported by the average genome-wide *F*ST estimated from all 11.5 million SNPs, which showed that genetic differentiation was much lower at Planoles (Pla*P*^m^ vs. Pla*S*^y^: *F*ST = 0.003) than it was at Avellanet (Ave*P*^m^ vs. Ave*S*^y^: *F*ST = 0.048) (Fig. 1E). In fact, *F*ST between Ave*P*^m^ and Ave*S*^y^ was higher than between Ave*P*^m^ and Pla*P*^m^ (*F*ST = 0.027), which are separated by more than 50 km, whilst Ave*S*^y^ and Pla*S*^y^ showed the highest pairwise *F*ST (= 0.066) of all.

The above results suggest a more substantial history of hybridization and gene flow at the Planoles hybrid zone than at Avellanet. To assess this more formally, we used the program *δaδi* (Gutenkunst et al., 2010) to fit a series of demographic models to the joint site frequency spectrum separately at each hybrid zone. We first compared the fit of a model of strict isolation (SI, where two populations diverge with no gene flow) to a model of secondary contact (SC, where populations diverge in allopatry followed by gene exchange after coming back into contact). For both hybrid zones, the SC model was a far better fit to the data than the SI model (ΔAIC > 2000 for both zones), providing evidence of gene flow between the magenta and yellow populations at each zone (Fig. S3, Table S2). It also suggested a more substantial history of gene flow at Planoles characterised by a much longer period since secondary contact than at Avallenet.

Together, these results suggest strikingly different histories of gene flow at each of the hybrid zones, which is largely consistent with observations made at these hybrid zones over more than a decade. At Planoles, plants are abundant every year, and hybrid individuals can be found over broad areas spanning more than 1 km (Whibley et al., 2006). In contrast, we do not always find a large number of plants at Avellanet (Stankowski, Barton & Field; personal observations). In some years, the plants are abundant, and in others, their distribution is patchy and hybrids are uncommon. Thus the difference in genetic structure between the zones may reflect the demographic stability of the populations which is what ultimately provides opportunities for hybridization and subsequent gene flow across the zone.

### 2.2 | Genome scans reveal highly heterogeneous differentiation landscapes with varying degrees of parallelism

Although we observed a strong difference in the magnitude of *F*ST at each hybrid zone, it is possible that finer-scale pattern of differentiation along the genome is highly similar. Indeed, highly correlated *F*ST landscapes have been observed in studies where multiple populations with varying levels of differentiation have been compared (Burri et al., 2015; Stankowski et al., 2019). The general explanation for observing correlated differentiation landscapes is that common evolutionary processes and intrinsic genomic properties have shaped variation across multiple incidences of divergence in isolation (Burri et al., 2015), local adaptation (Jones et al., 2012), or secondary contact (Nouhaud et al., 2022).

To test for correlated differentiation landscapes, we first calculated Hudson’s *F*ST in 10-kbp non-overlapping genomic windows for each pair of populations. This revealed highly variable patterns of differentiation among the comparisons, both in the level of *F*ST and pattern of heterogeneity. First, comparing the genome scans between var. *pseudomajus* and var. *striatum* at each of the hybrid zones, we found little heterogeneity in the pattern of differentiation at Planoles. *F*ST was consistently low across most of the genome (median = 0.008, sd = 0.011) (Fig. 2), with the exception of a small number of localised peaks of differentiation rising above the background. In contrast, the *F*ST landscape was highly heterogeneous at Avellanet (median = 0.028, sd = 0.064), with far greater variability across chromosomes and many areas with pronounced differentiation (Fig. 2). The windowed *F*ST estimates exhibited strong dissimilarity between the hybrid zones (Spearman’s rho = 0.04; hereafter, *ρ*) (Fig. S4A, Table S6).

**Figure 2.**
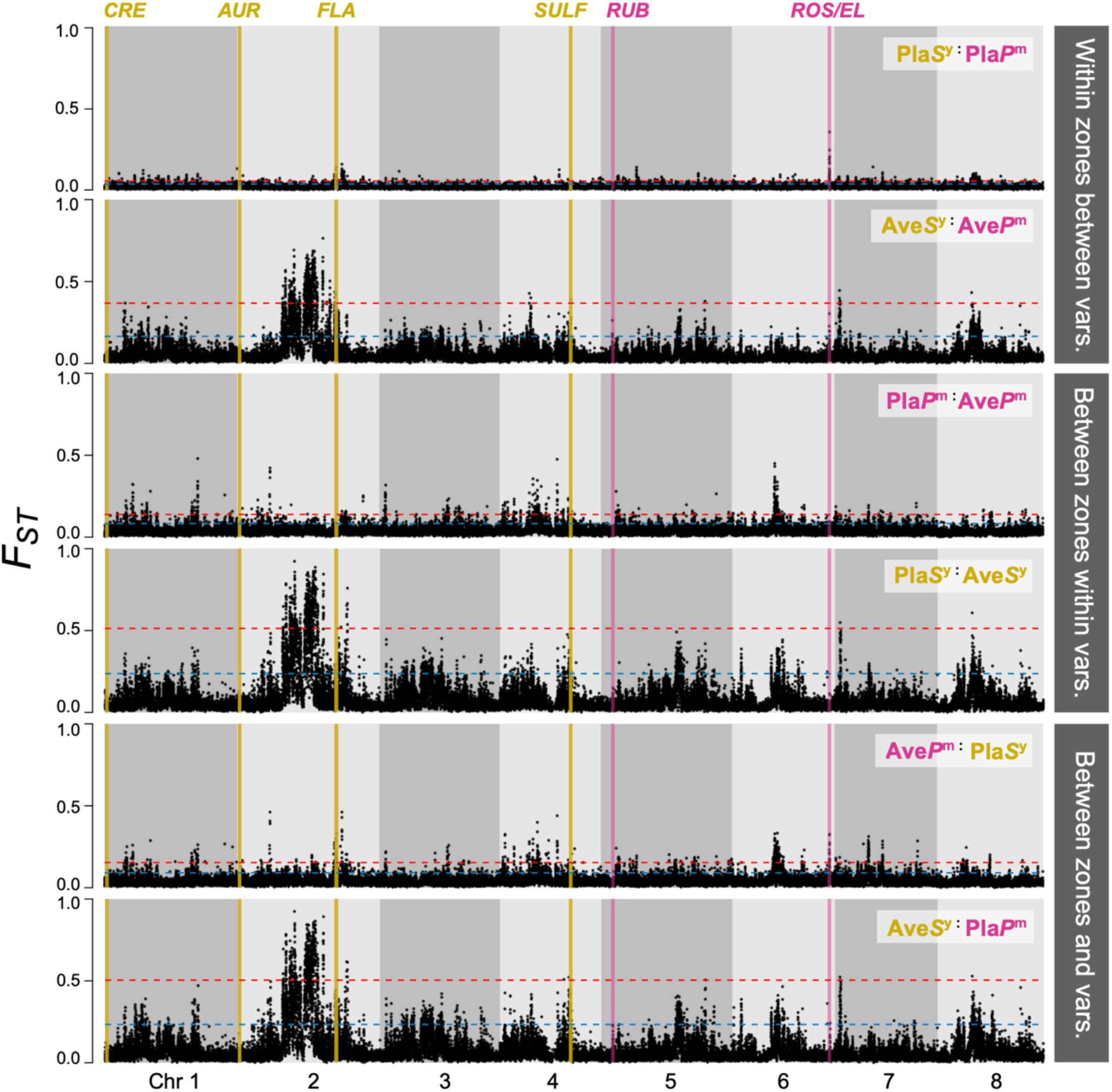
Genome scans show heterogenous *F*_ST_ landscapes with varying degrees of parallelism. *F*_ST_ is estimated in 10 kbp non-overlapping windows (*n* = 50,881) for each chromosome. Dotted blue and red lines show the 95th and 99th percentile of genome-wide *F*_ST_ estimates. **Top panel** (Within zones and between varieties): Comparison between varieties at each hybrid zone (Pla*P*^m^ vs Pla*S*^y^, Ave*P*^m^ vs Ave*S*^y^). **Middle panel** (Between zones and within varieties): Comparison between hybrid zones for magenta and yellow population (Pla*P*^m^ vs Ave*P*^m^ , Pla*S*^y^ vs Ave*S*^y^). **Bottom panel** (Between zones and varieties): Comparison between varieties from different hybrid zones (Pla*S*^y^ vs Ave*P*^m^, Pla*P*^m^ vs Ave*S*^y^). Grey shading delimits chromosomal boundaries. Pla: Planoles, Ave: Avellanet, *P*^m^: magenta-coloured var. *pseudomajus, S*^y^: yellow-coloured var. *striatum*.

The remaining comparisons showed that differentiation patterns depended heavily on the populations included. Most notably, Ave*S*^y^ showing highly parallel patterns (*ρ* ranging from 0.63 to 0.86) between comparisons that included it (Fig S4B, Table S6). This shows that the highly heterogeneous differentiation landscape at the Avellanet hybrid zone is driven more by the history of Ave*S*^y^ population than by gene flow between Ave*P*^m^ and Ave*S*^y^.

At both hybrid zones, elevated *F*ST windows tended to coincide with reduced genetic diversity (*πw*) in one of the two populations, and/or elevated between-population sequence divergence (*dxy*) (Fig. S5). At Planoles, the diversity landscapes were highly similar (*ρ* = 0.89), with outlier regions tending to show lower *πw* in Pla*S*^y^ (Fig S6A, Table S6). The diversity landscapes were less similar at Avellanet (*ρ* = 0.47), with the population that showed lower *πw* varying among the genomic regions (Table S6). For example, in the most pronounced *F*ST island on Chr 2, Ave*S*^y^ had lower *πw*, whereas Ave*P*^m^ had lower *πw* in the outlier regions on Chr 1 (Fig. S6B, S7). At Avellanet, we found a clear negative relationship between the local recombination rate and *F*ST (*ρ* = –0.30), indicating that highly differentiated regions at Avellanet tended to show lower recombination rates (Fig. S5, Table S6). Coupled with the low diversity and strongly elevated *dxy* in highly differentiated regions (Fig. S5), this suggests that *F*ST has been shaped primarily by widespread linked selection acting independently in the two populations. At Planoles, the relationship between recombination rate and *F*ST is less pronounced (*ρ* = 0.08), with outlier regions showing variable rates of recombination and modest reductions in *πw* (Fig. S5).

We next examined patterns of differentiation at known colour loci to determine whether they would be detected in outlier scans, since flower colour is expected to have evolved before the hybrid zone formations (Tavares et al., 2018). At Planoles, two of the known colour regions were identified as *F*ST outliers using both 95^th^ and 99^th^ percentile thresholds (Fig. 2, S4A). This included regions containing the *Flavia* locus (hereafter, *FLA*) on Chr 2 that controls the intensity of yellow pigmentation, and the two tightly linked loci *Rosea* and *Eluta* on Chr 6 (hereafter, *ROS/EL*) that have large effects on magenta colouration. At Avellanet, the *FLA* locus was identified at the 99^th^ percentile threshold, while *ROS/EL* was only detected at the 95^th^ percentile threshold (Fig. 2, S4A).

Overall, we found that the *F*ST landscapes at the two hybrid zones are quite different, which is not unexpected given their different demographic histories and patterns of hybridization. Although samples of var. *pseudomajus* and var. *striatum* from Avellanet are more physically distant than in Planoles (Fig. 1, Table S1), similarly distant samples at Planoles show a very similar *F*ST landscape to the one presented here (compare top panel in Fig. 2 to Fig. 5E in Field et al., 2025). While we were able to detect two of the major-effect colour loci at Planoles, we failed to detect the other known loci at either Planoles or Avellanet. This may be due to several factors. First, this dataset consists of low-coverage sequencing, so SNPs are sparser than a previously analysed high-coverage pool-seq datasets (Tavares et al., 2018). Second, previous work at Planoles has shown that the detection of smaller effect loci (e.g., *Rubia* and *Aurina*) depends on the proximity of sampling to the hybrid zone (Field et al., 2025). Our samples were collected very close to the point of contact, meaning that differentiation at small effect loci may have been swamped by gene flow. Finally, the large effect locus *Sulfurea* is a deletion polymorphism, so the functional allele may be more difficult to identify with SNP markers (Bradley et al., 2017).

### 2.3 | Different genealogical inference methods produce vastly different numbers of trees yet infer similar genealogical landscapes

Given the challenges of interpreting multiple pairwise *F*ST scans, we next shifted to genealogical tools that allowed us to jointly analyse relationships among all four populations. Among the variety of tools available, we selected and compared four that were broadly representative of the main approaches in methodological implementation: (*i*) Neighbour-joining trees in arbitrary windows (Martin & Van Belleghem, 2017), (*ii*) *tsinfer* (Kelleher et al., 2019), (*iii*) *Relate* (Speidel et al., 2019), and (*iv*) *Singer* (Deng et al., 2024). We restricted our comparison to a 2 Mbp region (45-65 Mbp in Chr 2) with 461,864 SNPs, that showed highly heterogenous patterns of *F*ST (Fig. 2) and contains the *FLA* locus (Bradley et al., 2025).

The first method divides the genome into non-overlapping windows containing the same number of SNPs (50 SNPs in our analysis) and infers a phylogenetic tree for each region separately. We inferred neighbour-joining trees, though other methods such as maximum-likelihood have also been applied (Fontaine et al., 2015). While simple and widely used, this approach has a significant limitation. Arbitrarily defined genomic segments often span historical recombination events, where relationships between haplotypes cannot (and ideally should not) be accurately represented by a single bifurcating tree (Shipilina et al., 2023). As a result, important genealogical signals may be dampened by the clumping of unique trees into one.

Unlike NJ trees, *tsinfer, Relate* and *Singer* infer a sequence of trees, consistent with how historical recombination events have altered genealogical relationships across the genome. They both do this by allowing topologies to vary locally to reconcile neighbouring site patterns that cannot be represented as a single bifurcating tree, but their approach varies significantly. *Tsinfer* reconstructs plausible ancestral sequences from sampled chromosomes and then infers the relationship between those sequences, preserving the correlation between consecutive genealogical trees. Therefore, neighbouring trees inevitably share many of the same nodes and branches. *Relate*, on the other hand, infers a completely new tree upon encountering an incompatible SNP. Therefore, consecutive trees do not share homologous nodes or branches, although this can be partly addressed by assigning the same age to nodes with identical descendant sets across adjacent trees.

Although *tsinfer* and *Relate* accommodate the effects of past recombination, they do not explicitly model recombination. In other words, incompatible SNP patterns only imply that historical recombination occurred somewhere between the boundary of neighbouring trees. *Singer* goes further and takes a Bayesian approach, attempting to infer the full ARG by fitting a model of coalescence and recombination to the SNP data. So, unlike the deterministic topology inference of *tsinfer* and *Relate* (i.e., multiple runs will always produce the same tree topologies, though inferred branch lengths may differ between runs), each MCMC iteration of *Singer* estimates a tree sequence that is drawn from the posterior distribution of possible trees. Thus, a region with no SNP may contain multiple inferred trees, that are not supported by any data within that region, but are a plausible outcome of the estimated model.

Examination of the resulting tree sequences shows that methods produce vastly different results (Table 1). First, we found that the number of trees varied substantially across methods. The neighbour-joining method contained the lowest number of trees at 9,237 (i.e, 1 tree for each 50 SNPs window). *Relate* and *tsinfer* inferred substantially more trees, with 198,375 and 406,135, respectively. *Singer* inferred the most trees by far, with 1,950,778. The average span of an NJ tree was 2.1 kbp (sd = 2.3 kbp), compared with 100 bp (sd = 322 bp) for *Relate*, 50 bp (sd = 222 bp) for *tsinfer*, and 9 bp (sd = 24 bp) for *Singer*. Finally, we asked how many SNPs fell within the span of each marginal tree. Marginal trees in *Relate* contained an average of 2.3 SNP (sd = 1.7) compared with 1.14 (sd = 0.47) for *tsinfer*. On average, trees for *Singer* contain less than 1 SNP (0.24, sd = 0.54), with 1950778 (80.73%) trees containing no SNP.

**Table 1.** Results of tree inference for four genealogical inference methods. The methods were applied to the same 2 Mbp region (45-65 Mbp in Chr 2) that contained 461,864 SNPs. The total number of trees inferred by each method, the mean span of trees in bp, and mean number of SNPs associated with each tree are provided.

| Inference method | Number of trees | Mean tree span (bp) | Mean no. SNPs per tree |
| --- | --- | --- | --- |
| Neighbour-joining trees in 50 SNP windows | 9,237 | 2126.44 | 50 |
| *Relate* | 198,375 | 99.81 | 2.33 |
| *Tsinfer* | 406,135 | 49.24 | 1.14 |
| *Singer* | 1,950,778 | 9.25 | 0.24 |

Overall, the characteristics of each tree sequence align with their respective methodological approaches. The number of SNPs associated with each NJ tree is defined by the user and will ultimately reflect a trade-off between information content (i.e., number of SNPs) and tree span. In an ideal world, window size would be minimised such that trees span as few recombination events as possible. However, if we assume that the transitions between trees by *tsinfer* reflect real recombination events, this would imply that the average 50 SNP window spans 43 observable recombination events. Although *Relate* and *tsinfer* define margins between trees based on incompatible SNP patterns, *Relate* produces half the number of trees. This may reflect different levels of tolerance for incompatibilities, and the number of trees may vary depending upon the parameters chosen by the user. Finally, because *Singer* models recombination explicitly, it inevitably produces far more trees than the other methods. Although many trees in the sequence are not supported by SNP data, allowing recombination to shape the sequence in the absence of polymorphism data is more consistent with reality, and may provide additional information in some inference schemes. However, it seems reasonable to exercise caution when making detailed inferences from trees that are not supported by SNP data.

Moving beyond the summaries of tree sequences, we next used topology weighting to compare how topologies change along the genome. Topology weighting iteratively subsamples one haplotype from each population and estimates the proportion of each subtree topology. In this dataset of four populations: Ave*P*^m^, Ave*S*^y^, Pla*P*^m^ and Pla*S*^y^, it weighs the contribution of each of the three possible topologies (Fig. 3) – the geography tree (Tgeo), where samples cluster by hybrid zone ((Ave*S*^y^,Ave*P*^m^)(Pla*S*^y^,Pla*P*^m^)), the variety topology (Tvar), where samples cluster by the variety, ((Ave*S*^y^,Pla*S*^y^)(Ave*P*^m^,Pla*P*^m^)), and an alternative topology (Talt), where samples neither cluster by geography or variety ((Ave*P*^m^,Pla*S*^y^)(Ave*S*^y^,Pla*P*^m^)). By iteratively sampling many subtrees (in our case 10,000), we can obtain their relative frequencies (i.e., topology weights), which provide a measure of the weight (or bias) of the full tree to each group level topology (Fig. 3).

**Figure 3.**
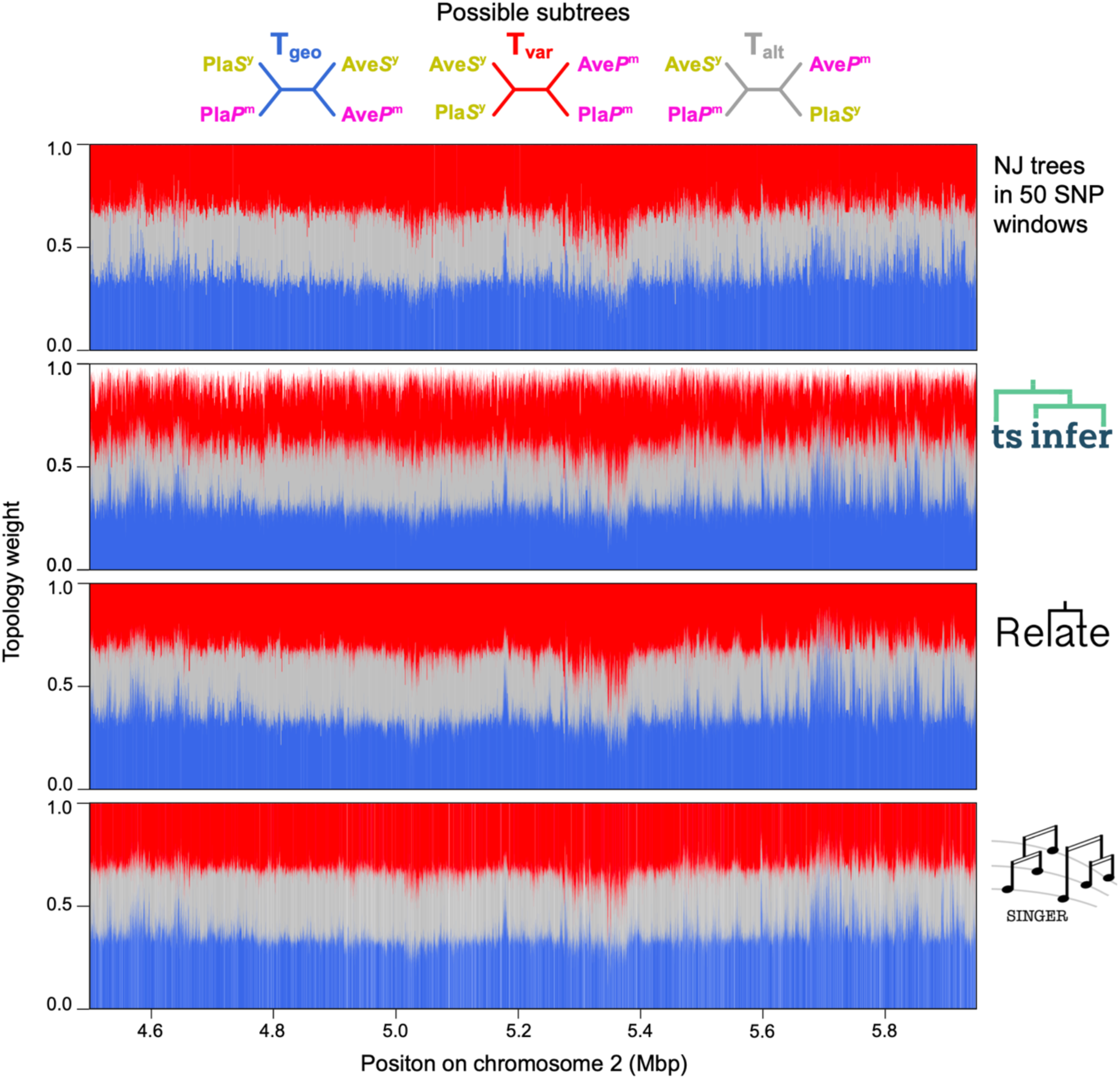
Topology weighting of trees sequences inferred by four different methods yield broadly similar genealogical landscapes. Topology weights of the three possible subtree topologies (T_geo_, T_var_, T_alt_) are plotted for each tree in the sequence along a small section of Chr 2. Each vertical bar shows the proportions of each topology in one genealogical tree. Therefore, topology weights add up to 1 except for trees inferred by *tsinfer* since it allows polytomies.

Although the characteristics of the tree sequences vary among the four methods, there is striking similarity in genomic distributions of the topology weights inferred from them. Figure 3 shows the weights for the three group-level topologies. From visual inspection alone, the topology weights are highly similar among the methods, increasing and decreasing in a coordinated way along the chromosome. Correlation analysis of the weights, performed on the topologies that coincide with SNP positions, shows the similarity is indeed quite strong among the methods (*ρ* value ranges for Tgeo: 0.57–0.77, Tvar: 0.57–0.66, Talt: 0.45– 0.71; Fig. S8, Table S7).

However, there are also some clear distinctions. First, the change in amplitude of the weights is not as extreme in the NJ method compared with the other methods. This is not surprising, as the 50 SNP windows span many distinct marginal trees, which we would expect to have a smoothing effect. Another major difference is that topology weights from *tsinfer* do not sum to one, implying that some subtrees cannot be classified as one of the 3 possible topologies. The reason for this is that *tsinfer* infers polytomies, while *Relate* and *Singer* force all branches to bifurcate.

In summary, the results of our comparisons show that the different methods produce vastly different tree sequences, yet largely agree on how group-level relationships change along the genome for our snapdragon dataset. It is good to know that the crudest and most sophisticated approaches give a similar picture, if only from a topological standpoint. Deciding which to use will depend on the size of the dataset and the goals of the study. However, we see little reason use the window-based approach given that more computationally efficient and precise methods are now available. *Tsinfer* and *Relate* are far better options but have different strengths. For example, *tsinfer* retains nodes and branches among trees, meaning that they can be represented as an ARG and used in analysis that leverage homology of tree features (Shipilina et al., 2023). In contrast, *Relate* has been shown to be more accurate than *tsinfer* (+*tsdate*) when it comes to estimating deeper coalescence times (Brandt et al., 2022). *Singer* is far more computationally demanding than the other methods, so it is difficult to scale to large datasets. However, for smaller datasets, and in a more defined genomic regions of interest, *Singer* allows for extremely fine-scale genealogical inference along with estimates of uncertainty.

### 2.4 | Topology weighting reveals regions associated with flower colour

For our purpose, *Relate* seemed to be the ideal choice to infer genome-wide genealogies, due to its scalability, efficiency, resolution of polytomies, and accuracy of inferring deeper coalescence times. Applying the algorithm to our genome-wide dataset yielded at sequence that contained 4,975,454 trees, with an average span of 101.2bp (sd = 739.4bp). We again used topology weighting to quantify bias toward the three group-level relationships (Tgeo, Tvar, Talt) for each marginal tree.

We first analysed the distribution of all topology weights in a ternary framework using the program *TwisstNtern* (Stankowski et al., 2024). The ternary plot is a natural framework for analysing the joint distribution of weights in a tree with four groups because it is possible to graphically represent each tree as a single point in an equilateral triangle based on the three weights. The three corners of the ternary plot—[1, 0, 0], [0, 1, 0], [0, 0, 1]— correspond to all of the trees where the sampled subtrees match only one of the three possible group-level subtrees. In contrast, the centre of the ternary plot—[1/3, 1/3, 1/3]—corresponds to where all three of the possible subtrees are found at equal frequency. Any other location in the ternary plot indicates an enrichment of one particular subtree topology. Previous simulations have shown that the ternary distribution of weights can be shaped by a range of factors, including population split times and effective population sizes, as well as processes that lead to haplotype sharing between non-sister groups (e.g., introgression) (Stankowski et al., 2024).

In our analysis, we expected the ternary distribution of weights to be skewed toward the geography topology (Tgeo, top of the triangle in Fig. 4), because this topology matches the genome-wide relationships observed between the populations (i.e., topology weighting of the genome-wide neighbour joining tree (Fig. S2) yields weights of Tgeo=1.0, Tc=0.0, Talt=0.0). While we did observe this skew, the bias toward Tgeo was relatively weak (mean Tgeo=0.367, Tvar=0.317, Talt=0.316), and similar across the 8 chromosomes. Although some of the trees showed high Tgeo weights (max Tgeo=0.93 with 5% of trees showing weights above 0.49), none of the 4,975,454 trees perfectly matched Tgeo. Rather, most of the genealogies clustered near the centre of the ternary plot (i.e., most weights were near 0.33 for all three topologies), indicating that haplotype diversity is broadly shared across the four groups.

**Figure 4.**
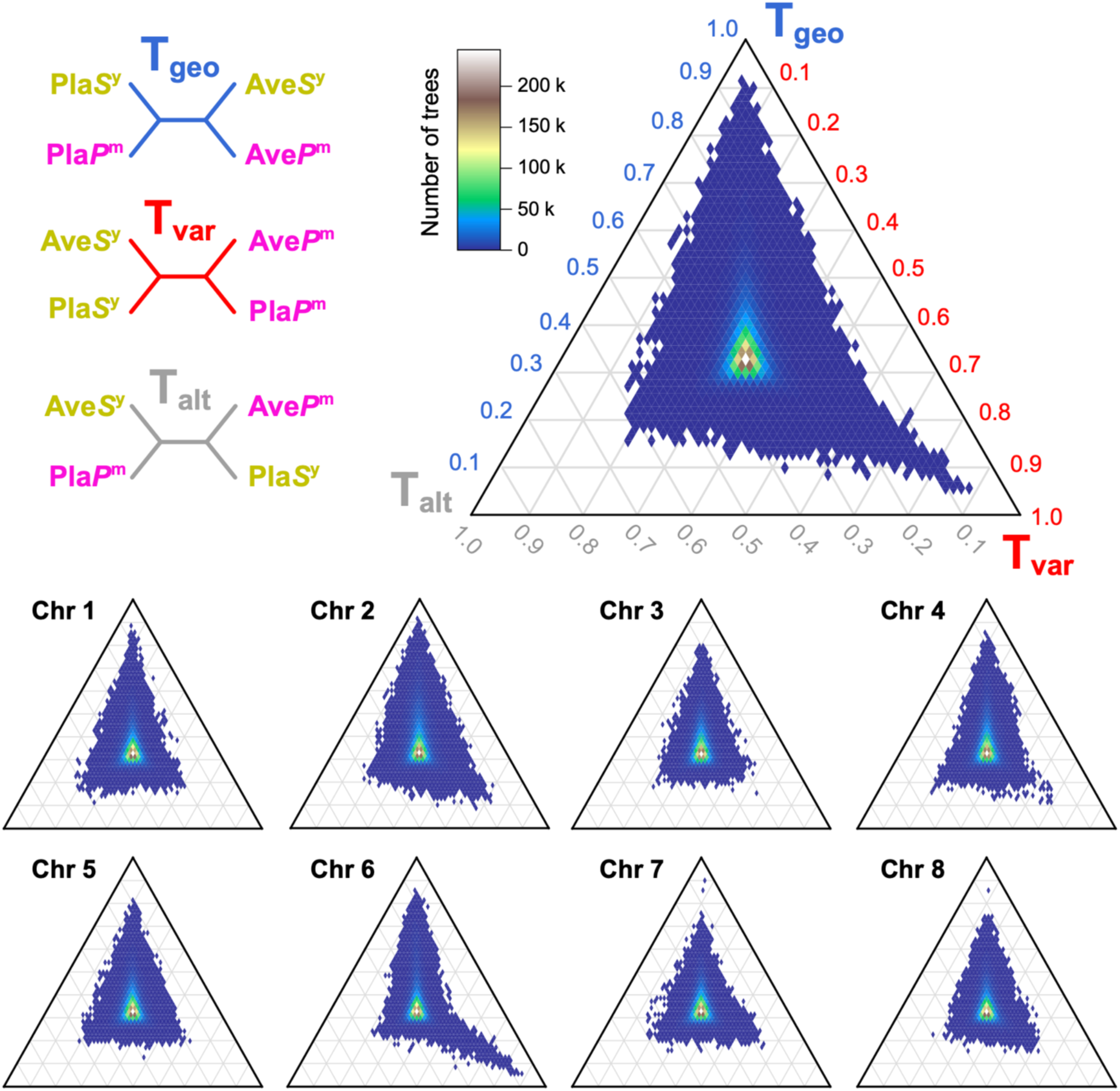
Ternary plots showing the joint distribution of topology weights. Empirical distributions of weights for the 4,975,454 trees inferred using *Relate* (top right), and for each chromosome (bottom). Each tile in the distribution is coloured according to the density of genealogies falling in that area of the distribution, as indicated in the colour scale. The three topologies associated with each axis in the ternary plot are shown in the top left of the plot. The three corners of the ternary plot— [1,0,0], [0,1,0], [0,0,1]—correspond to trees that perfectly match the three possible group-level subtrees.

Notably, we found striking left-right asymmetry in the distribution of topology weights between the left and right halves of the ternary plot (Fig. 4). Specifically, we observed a long tail of topology weights extending toward the right-hand corner of the plot, resulting in a 1% bias in the distribution toward the variety topology (Tvar). Such a bias is unexpected when sharing is due to the random sorting of ancestral polymorphism, as there is an equal chance that any given tree will be biased toward either one of the discordant topologies, leading to a symmetrical distribution of weights (Stankowski et al., 2024). This asymmetry is similar to what is measured by the site-based statistic Patterson’s *D*.

Indeed, roughly symmetrical distributions were observed on several chromosomes, including Chr 1, 3 and 5 (Fig. 4). The remaining chromosomes showed significant asymmetries toward the variety topology, driven by a relatively small number of genealogies (1,490 or 0.03%) with Tvar weights that exceeded 0.55 (Fig. S9). This indicates a bias of haplotype sharing between populations of the same variety (Fig. S9). The most striking bias was observed on Chr 6, where weights approached 0.9.

To explore regions associated with the genetic differentiation varieties, we plotted the genomic positions of detected Tvar outliers (Fig. 5). Tvar outliers were spread across multiple points along each of the chromosomes rather than clustering at a single site. Most of the known colour genes were observed near Tvar outliers, but we also observed bias toward Tvar in regions of the genome that have no known effect on flower colour, including regions of Chr 1, 5, 7 and 8.

**Figure 5.**
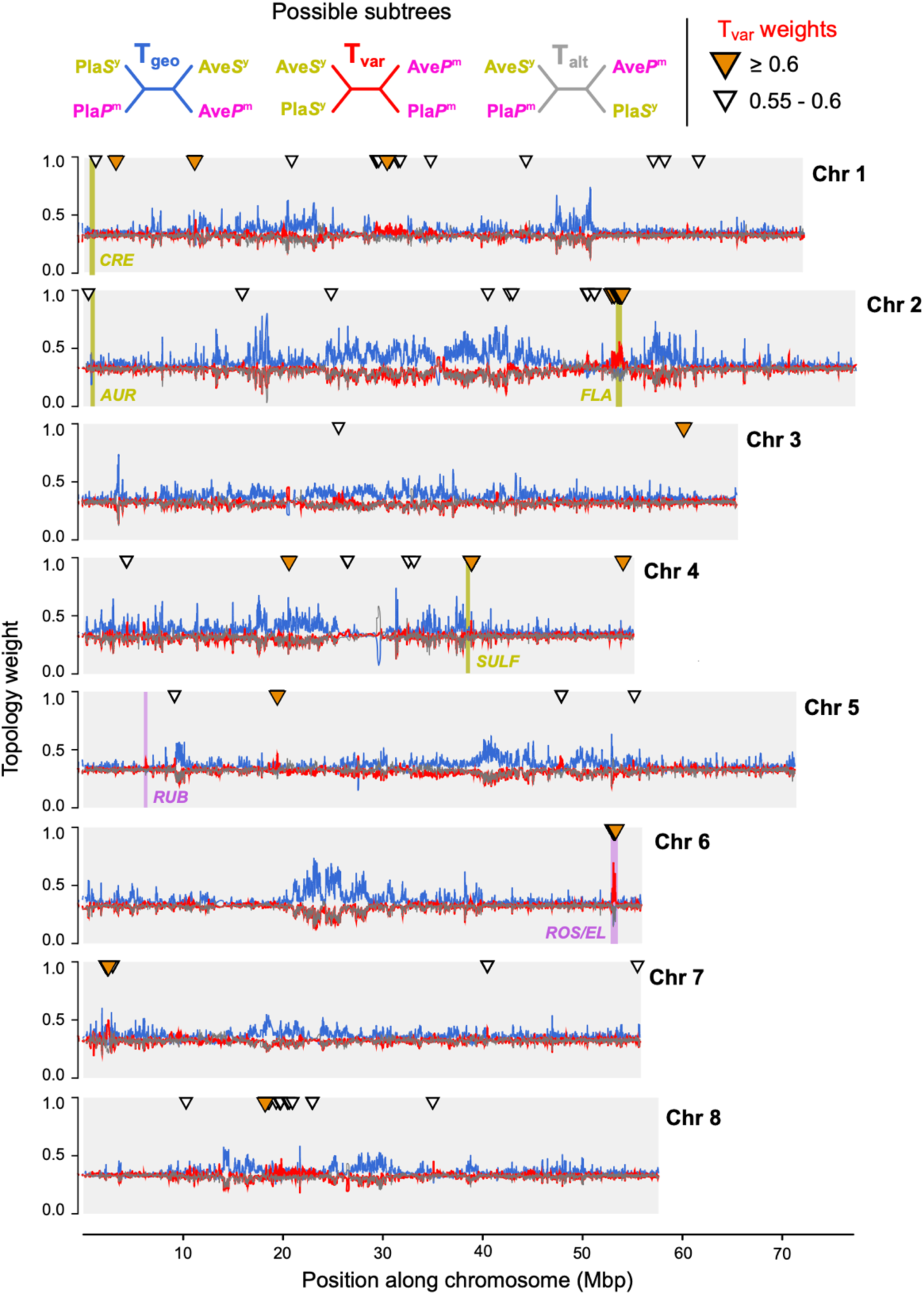
Genealogical landscape of parallel hybrid zone formation revealed by topology weighting. Topology weights (loess smoothed, span = 50 kbp) for the 4,975,454 trees inferred by *Relate* plotted along each chromosome. 7 loci controlling flower colour are highlighted in yellow or magenta. White triangles indicate trees with raw T_var_ weight between 0.55 and 0.6. Orange triangles indicate trees with raw Tvar weights ≥ 0.60.

### 2.5 | Coalescence times at FLA and ROS/EL differ from surrounding background

The two genomic regions that showed the clearest association with the colour topology were on Chr 2 and Chr 6, together accounting for 66% of all Tvar outliers (or 84% using the 0.6 cutoff). The outlier region on Chr 2 includes the recently discovered *Flavia* locus, which affects the patterning of yellow colouration in the face of the flower (Bradley et al., 2025) (Fig. 6). This signal of Tvar enrichment extends over roughly 2 Mbp of the chromosome, interrupting bias toward Tgeo on either side of it. Within the *FLA* locus, weights for some genealogies exceed 0.7 (Fig. 6). The other region, *ROS/EL* located on Chr 6 contains the two linked colour loci, *Rosea* and *Eluta*. *Rosea* activates anthocyanin biosynthesis across the corolla, while *Eluta* modifies its distribution (Tavares et al., 2018). Within the *ROS/EL* region, Tvar weights are strongly elevated and characterised by local peaks and troughs spanning about 1 MB. On either side of *ROS/EL*, all three topology weights hover around 0.33, indicating that haplotype variation is broadly distributed among the groups.

**Figure 6.**
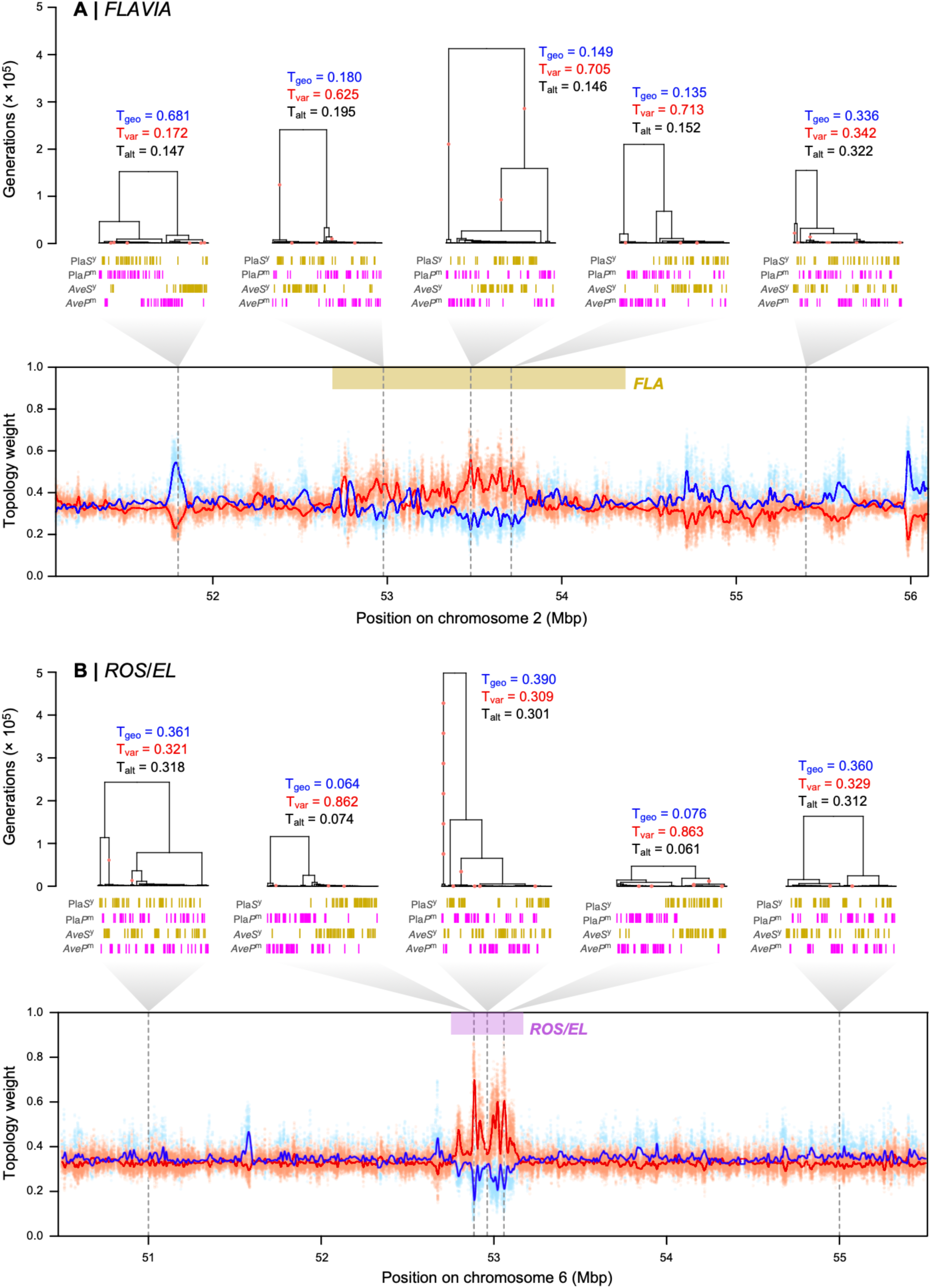
Fine-scale genealogical landscape at the *FLAVIA* and *ROS/EL* loci. **(A)** 5 Mbp genomic region centred around *Flavia*. Trees show relationships at various points along the sequence, with red circles indicating mutations associated with each tree. Vertical bars represent haplotypes, coloured according to the populations (Pla*S*^y^, Pla*P*^m^, Ave*S*^y^, Ave*P*^m^). From left to right (1-5), trees 1 and 5 are chosen arbitrarily, but equally distant from the locus. Trees 3 and 4 are trees have the highest smoothed and raw T_var_ weights, respectively. Bottom panel shows topology weights (T_geo_, blue; T_var_, red; T_alt_, black) through the region. Solid lines are loess smoothed weights (span = 50 kbp), while dots are raw weights. *FLA* locus (Chr2:52560000-54050000) is marked in a yellow bar, while the rest is considered as flank in TMRCA calculations. **(B)** Same as (A), but for *ROS/EL* locus. From left to right, trees 1 and 5 are equally distant from the colour locus, while trees 2-4 are within in. Tree 2 and 4 have the highest raw T_var_ weights at the *ROS1* and *EL* loci. Tree 3 shows a low T_var_ likely due to recombination between the 2 linked loci. *ROS/EL* locus (Chr6:52775000-53150000) is marked in a magenta bar, while the rest is considered as flank in TMRCA calculations.

Given existing evidence for selection on *FLA* (Bradley et al., 2025) and *ROS/EL* (Tavares et al., 2018), we next examined the coalescence times for genealogies in and around these colour loci. Since positive selection purges haplotype diversity from the population, we may expect to find shallower coalescence times within each variety (often measured using *π*w) reflecting the historical sweep of causal alleles (Hejase et al., 2020). In addition, these loci can also generate local barrier effects in the genome, which we would expect to increase coalescence times between the varieties (often measured using *dxy*) (Hejase et al., 2020; Wakeley, 2009).

To test for these patterns, we first compared the median time to the most recent common ancestor (TMRCA) for genealogies inside each locus to those in the flanking regions of the loci where there was no obvious association with colour (Fig. 6). For *FLA*, the median TMRCAs for var. *pseudomajus* (i.e., Pla*P*^m^+Ave*P*^m^) was higher inside the locus than in the flanking regions, whereas, for *ROS/EL*, there was no obvious difference (Fig. 7). For var. *striatum* (i.e., Pla*S*^y^+Ave*S*^y^), we observed a similar result in *FLA* and *ROS/EL*, with median TMRCAs being lower in the loci than in the flanking regions. We also compared TMRCAs between the varieties, finding higher median TMRCAs inside *FLA* and *ROS/EL* loci in comparison to the flanking regions.

**Figure 7.**
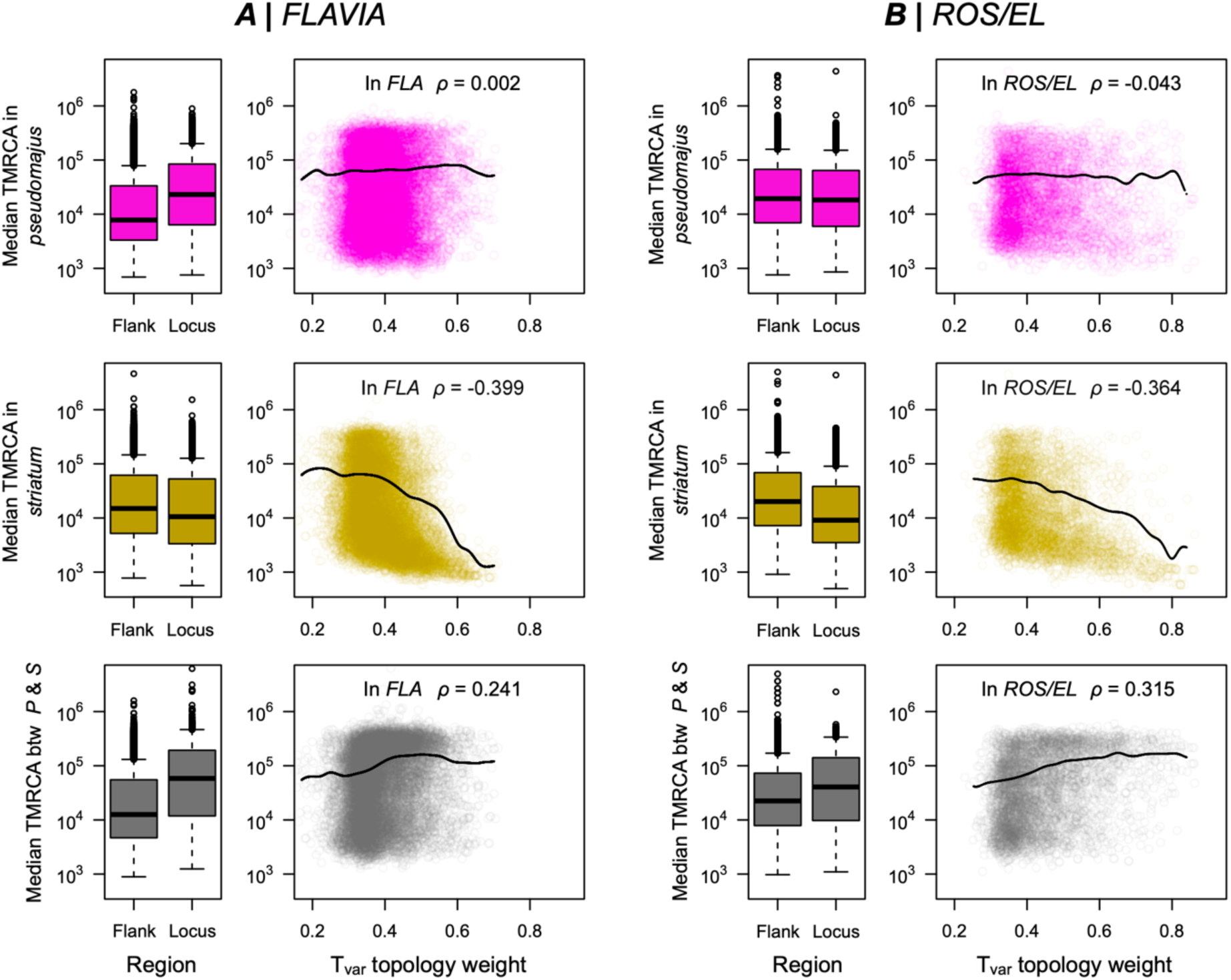
Coalescence times at *FLAVIA* and *ROS/EL*. Boxplots for (A) *FLAVIA* and (B) *ROS/EL* show the median time to most recent common ancestor (TMRCA) within var. *pseudomajus* (top row), within var. *striatum* (middle row), and between var. *pseudomajus* and var. *striatum* (bottom row), all on a log scale. The left boxes (‘Flank’) show the TMRCAs in the flanking regions around the locus, while the right boxes (‘Locus’) show the values inside the locus. The scatterplots and dashed black lines show the full distribution and the overall smoothed trend between the T_var_ weight and median TMRCA within the locus, (rows are as indicated for the boxplots). *ρ* is the correlation coefficient from a Spearman’s rank correlation.

We also examined the relationship between the TMRCAs and Tvar weights inside each locus, as we expected signatures to be most pronounced for genealogies that more closely resembled the variety topology. In *FLA*, we found no relationship between the median TMRCA and Tvar weight within var. *pseudomajus* (*ρ* = 0.002, Fig. 7). For *ROS/EL*, we only observed a weak negative relationship, with higher Tvar trees showing a broad range of median TMRCAs within var. *pseudomajus* (Fig. 7). In contrast, we found a very clear negative relationship between median TMRCA and Tvar weight within var. *striatum* for both loci, such that genealogies with high Tvar weights tended to have shallower median TMRCAs (*ρ* = –0.399 for *FLA* and –0.364 for *ROS/EL*, Fig. 7). We also observed a clear positive relationship between Tvar weights and median TMRCAs between the two varieties, such that genealogies with a high Tvar tended to have higher TMRCA or deeper coalescence times. Similar patterns are observed when each hybrid zone was analysed separately (Fig. S10, S11).

While preliminary, these results are consistent with (i) selection having acted on haplotypes associated with colour and (ii) suggests that these loci have a local barrier effect. For *FLA*, evidence for selection mainly comes from sharp allele frequency clines at the Planoles hybrid zone (Bradley et al., 2025; Field et al., 2025). Our results provide preliminary evidence that a selective sweep has occurred on the background of var. *striatum* and the allele is present in both Avellanet and Planoles. The lack of a signature in var. *pseudomajus* is consistent with the phenotypic effect of *FLA*, as it only affects yellow colouration. The *ROS/EL* locus is more complex, with previous evidence suggesting that there have been multiple independent sweeps at the two linked loci (Tavares et al., 2018). Here, we only found evidence for selection in the yellow group, while previous work has suggested sweeps on both the yellow and magenta backgrounds. Our results do not preclude such a sweep, as footprints of selection are transient and fade with time. However, it might indicate a more recent sweep in the yellow population. Also, our analysis is coarse-grained and does not consider how fine-scale genealogical relationships change across the region. In future, we plan to use genealogical tools and a much larger dataset to dissect this region in fine detail.

### 2.6 | Conclusion and implications for genomic studies of speciation

In this paper, we studied the genomic landscape associated with replicate hybrid zone in *Antirrhinum majus*. Our study highlights many of the known challenges in interpreting genome scans in the context of adaptation and speciation, as differentiation landscapes can be shaped by a multitude of factors and processes that have nothing to do with speciation *per se* (Ravinet et al., 2017; Wolf & Ellegren, 2017). Comparing two parallel hybrid zones, we show that genome scans can be dominated by signals of historical demography, a factor less widely discussed but critical for isolating speciation related patterns. At Planoles, var. *pseudomajus* and var. *striatum* show very little differentiation, as we would expect between taxa that were described as varieties based only a difference in flower colour. At a second previously unstudied hybrid zone at Avallenet, differentiation was far more striking and characteristic of more divergent taxa (Bolnick et al., 2023; Stankowski & Ravinet, 2021). This shows that differentiation landscapes can be extremely variable within species, highlighting the dangers of generalising about broader processes from a single pair of samples.

Our analyses suggest that different levels of divergence at the two hybrid zones are primarily due to variation in the timing and/or rate of gene flow following secondary contact. This raises important questions about what has caused this difference, and, more broadly, the long-term dynamics of gene flow between *Antirrhinum* varieties in the Pyrenees. The strongest patterns of differentiation were always observed in comparisons that included var. *striatum* from Avellanet. This result can be explained in several ways. For example, varieties at Avellanet may exhibit historical divergence that was typical of allopatric var. *striatum* and var. *pseudomajus.* This divergence may have been maintained at Avellanet, either by a lower rate of gene flow than at Planoles, or by secondary contact being much more recent than at Planoles. The current spatial distribution of *A. majus ssp. pseudomajus* hints at the first possibility, as the continuous populations around Planoles contrast with the patchier distribution at Avellanet. Moreover, a slightly distant population of *var. striatum,* isolated from Planoles by a mountain pass, shows a similar pattern of differentiation when compared to populations at Planoles (Field et al., 2025). Another possibility is that the var. *striatum* population from Avellanet were historically isolated, causing its demographic history to be distinct from other populations. These questions are beyond the scope of our current dataset, and more detailed work is needed to understand the biogeographic and evolutionary history of *A. majus*.

Topology weighting of marginal trees inferred from genealogies allowed us to identify loci associated with the two snapdragon varieties. Flower colour is the only trait that consistently differs between the varieties, suggesting that the Tvar outliers may underpin this variation. Two of the loci known to cause differences in pigmentation, *Flavia* (Bradley et al., 2025) and *Rosea/Eluta* (Tavares et al., 2018), were identified as Tvar outliers; whilst others, *Cremosa* (Richardson et al., 2025), *Rubia* (Field et al., 2025), *Sufurea* (Bradley et al., 2017) and *Aurina* (Richardson et al., 2025) did not show clear associations. This could be due to several reasons, including proximity of samples to the core of the hybrid zones, effect size of the loci and sequencing coverage. Most of the Tvar outliers have not been previously associated with colour. It is possible that these outliers underpin some other trait (floral or non-floral) that differs between the varieties, or they may simply be spurious associations reflecting the highly stochastic nature of the coalescent process. This highlights a more general limitation of *all* genome scans: they detect regions of elevated differentiation between populations, and more detailed mapping studies and functional work are needed to demonstrate causality.

Finally, we encourage others to explore and critically evaluate the utility of genealogical methods in their research. Several recent studies, mostly focusing on human populations, suggest that genealogical tools can lead to more accurate inferences about past evolutionary processes (Fan et al., 2022, 2023; Speidel et al., 2019; Stern et al., 2019; Wohns et al., 2022). However, relatively few studies have used genealogical methods to study adaptation and speciation (Campagna et al., 2017; Hejase et al., 2020, 2022; Hooper et al., 2024; Meyer et al., 2024; Rueda-M et al., 2024; Stankowski et al., 2024; Wang & Coop, 2022). Our preliminary genealogical comparison of known adaptive loci with the surrounding genomic background further highlights the potential of these tools for studying the interplay between selective sweeps and barriers to gene flow. ARGs and tree sequences are very rich structures that are complex and challenging to interpret (Shipilina et al., 2023). However, paired with new linked-read sequencing methods, we think there is tremendous scope for creativity around how we can best visualise local genealogical relationships, account for uncertainty, and identify signatures that are associated with the speciation process.

## 3 | Materials and Methods

### 3.1 | Sample collection, DNA extraction, and sequencing

Leaf material was collected from individuals of *A.majus* at two hybrid zones near the towns of Planoles (42.3162°N, 2.1039°E) and Avellanet (42.3503°N, 1.3288°E) (Table S1). Several leaves were collected from each individual and refrigerated at 4°C before further processing. DNA was preserved by placing leaf tissue in a paper envelope, and envelopes into an air-tight plastic bag with silica gel. DNA was extracted using a custom protocol optimised for isolating high molecular weight DNA (Supplementary Methods).

### 3.2 | Library preparation and sequencing

Sequencing libraries were constructed by mixing genomic DNA with a pool of haplotagging beads with a different set of A and C barcode oligos (see Supplementary Table 1 for oligonucleotide sequences). This modification shifts the barcode position for the A/C segment from the original i7 index position into Read 2, followed by a mutated Tn5-17A/18G-MEmut sequence – ACTTGTGTATAAGAGACAG (Steiniger-White et al., 2002). The mutated Tn5-MEmut sequence allows tagmentation but does not otherwise interfere with Illumina sequencing. An additional standard 8-bp i7 Illumina index barcode was added during the final PCR amplification to introduce a fifth barcode segment to allow multiplexing of more than 384 samples. Amplified libraries were cleaned up and size-selected using Ampure magnetic beads (Beckman Coulter), Qubit quantified, and adjusted with 10 mM Tris, pH 8, 0.1 mM EDTA to 2.5 nM concentration for sequencing. Libraries were sequenced aiming for 2x coverage with Illumina paired-end sequencing (2x 150 bp) across a lane of Novaseq 6000 S4 by Azenta Life Sciences (Leipzig Germany). The sequences were then demultiplexed by recognising and trimming away the Tn5-MEmut sequence from R2 and the remaining B/D and A/C along with the Plate barcodes. The remaining sequences were processed as previously described in Meier et al. 2021

### 3.3 | Processing of raw reads and read mapping

Raw reads were mapped to the *A. majus* reference genome v3.5 (M. Li et al., 2019) using *EMA v0.7.0* (Shajii et al., 2018), a BX-tag-aware modification of *BWA* (H. Li, 2013). First, haplotag barcodes with BX tags were converted to 16-basepair barcodes using 16BaseBCGen (https://tinyurl.com/SamHaplotag). Reads with correct BX-tags (98.14%) were then mapped with *EMA*, which favours alignments where reads with the same barcode group together. Reads with faulty BX-tags (1.86%) were mapped to the genome using *BWA v0.7.17*. The resulting BAM files were combined and checked for quality using the *multi-bamqc* command in *qualimap v2.2.1* (Okonechnikov et al., 2016) (Table S1). PCR and optical duplicates were marked and removed using the *markdup* tool in *sambamba* (Tarasov et al., 2015).

### 3.4 | Variant discovery imputation, phasing and allele polarization

We used the *mpileup* and *call* commands in *bcftools v1.18* (Danecek et al., 2021) to identify candidate sites that were then used for final genotype inference and imputation by *STITCH v1.6.10* (Davies et al., 2016). Variant calling was performed with the *bcftools* multiallelic calling program using the flags *-m* and *--annotate AD,ADF,ADR,DP,QS,SP*. The resulting VCF was filtered to remove low-quality and potentially erroneous variant sites (Table S2). We first removed all INDELs (*bcftools view -V indels*), all SNPs within 5 basepairs of INDELs (*bcftools filter –SnpGap 5*), all monomorphic REF or ALT sites (*bcftools view -m2 - e “AC==AN || AC==0*), and all sites with more than 2 alleles (*bcftools view -M2*). Next, we removed all sites with >2.5 times the mean coverage across all samples (130x), sites with a genotype quality score <20 and a mapping quality score <30 (*bcftools filter -e “INFO/DP>130 | QUAL<20 | MQ<30*). Finally, bi-alleleic sites with >0.8 of missing genotypes were removed (*bcftools view -e “F_MISSING>0.80”*), producing a set 11,574,426 candidate sites (Table S2).

We applied *STITCH* to impute variants for the 11 million sites described above. *STITCH* models each chromosome as a mosaic of *K* founding haplotypes using both the underlying sequence reads and the linked-read information encoded in the BX-tag. Unlike traditional callers, *STITCH* imputes genotypes in the presence of missing data based on haplotype information from all sequenced individuals. Following guidelines and informed by pilot *STITCH* runs, we used the following parameters: *--K=75, --nGen=100, --niter=40, -- expRate=0.5, --downsampleToCov 10 --use_bx_tag TRUE*. To optimise computational resources and runtime, we performed *STITCH* with the above parameters on 1 Mbp regions with an overlap of 100kb overhang allowing them to be combined afterwards. Out of the 11 million sites, 41,396 (0.4%) sites were deemed invariant by *STITCH* (*i.e.,* the *bcftools* and *STITCH* calls disagreed) and were removed leaving a final set of 11,533,030 SNPs of which 93.9% had an INFO score ≥ 0.8, computed by *STITCH* as a proxy for imputation confidence (Table S3). Moreover, the observed and imputed allele frequency were highly correlated (*R^2^* = 0.87).

Finally, we used the *phase_common_static* from *SHAPEIT5 v 5.1.1* (Hofmeister et al., 2023) to statistically phase genotypes without a reference panel. We polarised alleles in *A. majus* as ancestral or derived using high-coverage PoolSeq sequence data (mean coverage = 89.97x) from multiple populations of the closely related outgroup species *A. molle* (Durán-Castillo et al., 2022). Detailed information on the logic used can be found in the supplementary methods.

### 3.5 | Genome-wide evolutionary relationships and demographic inference

We used three methods to infer genome-wide evolutionary relationships among the sequenced samples. First, we estimated principal components of the genotype matrix. Prior to analysis, we pruned the dataset to reduce linkage disequilibrium (LD) between neighbouring SNPs (*r*^2^ threshold of 0.1, window size = 50 SNPs, step size = 10 SNPs). This was done using *Plink v2.0 (Chang et al., 2015)* using the command *--indep-pairwise 50 10 0.1*, yielding 1,710,010 SNPs.

We used the model-based clustering program *Admixture v1.3* (Alexander et al., 2009), to assess the genetic structure. *Plink v2.0* was first used to produce BED files from the original VCF file. We ran *Admixture* on the LD-pruned dataset using the unsupervised model for all values of *K* ranging from 2 to 6.

We also inferred a phylogenetic network using the R package *phangorn v2.12* (Schliep, 2011). The LD-pruned dataset was converted to PHYLIP format using the script *vcf2phylip*. We then calculated a distance matrix from all aligned SNPs using the *dist.ml* function with *model = “JC69“*. The phylogenetic network was then inferred using the *neighborNet* function and drawn with *Splitstree v4.19.1* (Huson & Bryant, 2006).

We calculated per-site *F*ST between each pair of populations on the full SNP dataset, using the approach described by Weir and Cockerham (1984), implemented in *vcftools v0.1.16* (Danecek et al., 2011) using the *--weir-fst-pop* flag. Site-based estimates were averaged to obtain a genome-wide estimate.

We estimated gene flow between the two varieties independently in each locality using diffusion approximation, implemented in the software program *δaδi* (Gutenkunst et al., 2010). Since secondary contact is considered the most likely explanation for the current distribution of the two varieties (Tavares et al., 2018), we focused on comparing secondary contact models (SC) with strict isolation models (SI). All models included variation in the ancestral population’s effective size prior to population split, following Momigliano et al. (2021). In their basic form, both SI and SC models represent a population split into two populations with specific effective population sizes (N1 and N2) that diverge for a period without gene flow (Ts). In the SC model, these populations then begin exchanging migrants during a secondary contact phase (Tsc), with potentially asymmetric migration (M1 and M2). We expanded these models to account for recent population growth (p1 and p2) and/or Hill-Robertson interference by fitting a genome fraction (P) where the effective population is only a fraction (hrf) of what is found in the rest of the genome. In total, we tested 8 distinct models, including 4 modifications of the SI and SC models: (1) standard model (2) model with population growth in the daughter populations (3) a standard model with Hill-Robertson interference and (4) a combined model that included both population growth and Hill-Robertson interference. Each model was fitted 30 times to the data to ensure convergence, and model comparison was performed using the Akaike Information Criterion (AIC). The importance of gene flow in each locality was then compared by calculating the ratio between Tsc/Ts.

### 3.6 | Genome-wide differentiation, diversity, and recombination rate

We calculated *F*ST in 10 kb windows for each pair of populations using the script *popgenWindows.py* (https://github.com/simonhmartin/genomics_general). Genetic diversity was measured for each site using the *-site-pi* function in *vcftools*.

We used *LDhat v2.2* (Auton & McVean, 2007) to calculate the population-scaled recombination rate (*ρ*) between each SNP, separately for each population. We first used the *lkgen* function in *LDhat* to generate a log-likelihood lookup table for the number of haplotypes in each population, with *θ* = 0.009 (calculated from average genome-wide π in 10kb windows from previously published study (Tavares et al., 2018). We then used the *interval* function with the parameters: *-its 10000000 -samp 5000 -bpen 5*, to estimate variable recombination rates. Finally, we summarised results from the MCMC iterations to estimate mean *ρ* between each SNP using the *stat* function with the parameters: *--burnin 1000*. *LDhat* was performed on windows of 2000 variants with an overlap of 100 variants at each end and combined afterwards.

### 3.7 | Genealogical inference

We used four methods to infer trees from our data. First, we inferred neighbour-joining trees for 50 SNP non-overlapping windows using the script *phyml_sliding_windows py* (https://github.com/simonhmartin/genomics_general) with *–minPerInd = 15*.

The second method used was *tsinfer* (Kelleher et al., 2019). We used a custom script to convert phased, polarised SNPs into the *tskit.samples* format, which was then used to infer tree topologies with the *tsinfer.infer* function in *tsinfer v0.3.2* library (https://github.com/tskit-dev/tsinfer), followed by the *TreeSequence.simplify* function in *tskit v0.5.8* library (https://github.com/tskit-dev/tskit) to remove unary nodes.

Third, we inferred a tree sequence using *Relate v1.1.8* (Speidel et al., 2019). We assumed *μ* = 5.7 ✕ 10^-9^/bp/generation and uniform recombination rate of 1cM. We initially ran Relate separately on each chromosome setting the haploid *Ne* to 813388, as derived earlier from *π* = 4*Neμ* where *π* = 0.009. We then used the *EstimatePopulationSize.sh* script to jointly infer a time-varying population size history and branch lengths under that history. For this step, we used a *--threshold 0* to ensure that no trees were excluded in the joint-fitting and *--num_iter = 10*. We also included each population in the argument. Finally, we converted the genealogical trees stored in .*anc* and .*mut* format to .*newick* format with the *RelateExtract – mode AncToNewick* function. We focused our analysis on a 5 Mbp region around two flower colour loci: *FLAVIA* (locus – Chr2:52650000-54050000; region including locus and the flanking sequence on either side – Chr2:51100000-56100000) and *ROS/EL* (locus – Chr6:52775000-53150000; region including locus and the flanking sequence on either side – Chr2:50500000-55500000). Specific genealogical trees were plotted using a custom script modified from *Treeview.sh* in *Relate* library. Time to the most recent ancestor (TMRCA) are computed using a custom modified script from *tskit* library for 3 subsampled groups: within all var. *pseudomajus* individuals, within all var. *striatum* individuals and between var. *pseduomajus* and var. *striatum* individuals. For each case, TMRCA is first computed for all pairwise combinations of individuals, followed by calculating the median.

Finally, we ran *Singer v0.1.7* (Deng et al., 2024) on 500 kbp genomic windows. For each window, we calculated average *π* with *VCFtools*, which was then used to calculate *Ne* from *π* = 4*Neμ*. We ran *singer_master* with the parameters: *-m = 5.7e-9, -ratio = 1, - mcmc_iter = 100, -thin = 20, -polar = 0.9.* We then used the function *convert_to_tskit* to convert the last MCMC iteration to *tskit* format and to extract trees in *newick* format.

### 3.8 | Topology weighting and ternary analysis

Topology weighting was performed on sequences of trees derived from the various genealogical inference methods using *Twisst* (Martin & Van Belleghem, 2017). Due to the large number of trees and haplotypes, we followed standard *Twisst* guidelines and limited the topology sampling to 10,000 subtrees using the flag *--method fixed*. Genome-wide topology weights were plotted with loess smoothing (span = 50 kbp). We used the *TwisstNTern* framework (Stankowski et al., 2024) to visualise and calculate asymmetry in the distribution of topology weights for the whole genome, and for each chromosome separately, using the *--superfine* granularity.

## Author Contributions

Conceptualization: AP, DLF, DS, GC, NHB, SS, YFC; Formal Analyses: ALM, AP, DS, SS, YFC; Funding Acquisition: DLF, NHB; Resources: AJM, JKG, MK, NHB, YFC; Supervision: GC, NHB, SS; Visualization: AP, SS; Writing – Original Draft Preparation: AP, SS; Writing – Review & Editing: ALM, AP, DLF, DS, GC, NHB, SS, YFC. Data Curation: AP.

## Acknowledgements

We thank ESEB Godfrey Hewitt Mobility Award and Graham Coop for supporting AP’s research stay at UC Davis. We thank Tom Ellis, Parvathy Surendranadh and other Barton Group and Coop Lab members for stimulating discussions. We are grateful to all the interns and volunteers, who have helped us with fieldwork. We thank Eva Salmerón Mateu for her assistance in fieldwork logistics at the field station, El Serrat. We are grateful to Enrico Coen and his research group for providing the *Antirrhinum molle* PoolSeq data, used in the allele polarization. We are also thankful to Enrico Coen and Cristophe Thébaud for discovering the Avellanet hybrid zone, followed up with sampling led by DLF in 2017. The study was supported by Austrian Science Fund (FWF) Grant (Snapdragon Speciation P32166, awarded to DLF); ERC (Advanced Grant HaplotypeStructure 101055327, awarded to NHB); ERC (POC Grant 101069216, awarded to YFC) and the National Institutes of Health (NIH R35 GM136290, awarded to GC). YFC was supported by the Max Planck Society. Computing infrastructure for bioinformatics and analyses was provided by ISTA High Performance Cluster.

## Data Accessibility and Benefit-Sharing

Genomic sequence data produced and analysed in this study are deposited at the European Nucleotide Archive (ENA) under the BioProject accession number PRJEB88592. Original code for analyses can be accessed at Github (https://github.com/arka-pal/snap_replicateHZ). All samples were collected in accordance with the Nagoya Protocol on Access and Benefit sharing (Permit No. 170512ESNC3/ABSCH-IRCC-ES-237652-1). Benefits from this research accrue from the sharing of our data and results on public databases as described above.

