## Supplementary Materials for "Genealogical analysis of replicate flower colour hybrid zones in *Antirrhinum*"

This document includes:

- Supplementary Methods
- Figures S1 to S11
- Tables S1 to S7

### 1 | Supplementary Methods

#### *1.1 | DNA extraction*

DNA was extracted from a 1x0.5 cm$^2$ piece of dry leaf using a custom protocol optimised for isolating high molecular weight DNA. *(i)* Tissue was first crushed to a fine powder in a 1.5 ml Eppendorf tube using a micropestle. *(ii)* 0.5 ml of lysis buffer was added, containing 400 µl PureLink Genomic Digestion Buffer (Thermofisher: K182301), 40 µl Proteinase K (20mg/ml), 60 µl of 18% Polyvinylpyrrolidone (40 kdaG), and 2% Beta-mercaptoethanol. *(iii)* Samples were incubated for 15-20 mins at 60ºC with mixing at 950 rpm with occasional inverting of the tube, followed by 30 secs on ice. *(iv)* 10 µl of 5 mg/ml RNaseA was added and mixed by inverting the tube several times, and incubated for 5 mins at room temperature. *(v)* 155 µl 5M of potassium acetate was added and immediately mixed by inverting 10-15 times, followed by incubation for 2 mins on ice. *(vi)* The samples were centrifuged at 13k g for 10 mins at 14ºC and 400 µl of lysate was transferred to a new tube using a wide-orifice tip. *(vii)* 200 µl magnetic beads were added and immediately mixed by inverting 10-15 times and incubated for 10 mins at room temperature. *(viii)* Samples were pulse-spun for 2 seconds, and put on a magnet stand for 5 mins and the lysate was discarded. *(ix)* The beads were washed twice for 1 min with 80% EtOH; All EtOH was carefully removed after the second wash and the sample left open to evaporate for 2-4 mins at room temperature. *(x)* Samples were removed from the magnet and 100 µl of 10 mM Tris pH=8, 0.2mM EDTA was added to the tube and incubated for 10 mins at 42ºC to elute the DNA. *(xi)* Samples were mixed gently by inverting to re-suspend the beads and allowed to incubate for 10 mins at 42ºC. Eluted DNA was stored with beads at 4ºC. The concentration of each DNA sample was determined using a Qubit fluorometer or an M200 Pro Tecan plate reader. A rough estimate of size and quality was determined using agarose gel electrophoresis. Samples were diluted to 5 ng/µl using Tris, pH=8, 0.2mM EDTA and stored at 4ºC for future use.

#### *1.2 | Polarising allele in A. majus as ancestral or derived*

We polarised alleles in *A. majus* as ancestral or derived using high-coverage PoolSeq sequence data (mean coverage = 89.97x) from multiple populations of a closely related outgroup species *A. molle*. For each bi-allellic site in *A. majus*, we determined which alleles were present within *A. molle*. For 83.64% of sites, we observed 3 site-patterns that allowed us to identify the derived allele in *A. majus* (Table S4)

**Pattern A:** *6,624,396 sites (57.4%) resolved.* Sites where one of the *A. majus* alleles is fixed in *A. molle*. Out of these 6.6 million sites that constituted 57.44% of all the bi-allelic *A. majus* sites, 5.5 million (47.83%) sites had the *A. majus* major allele fixed in *A. molle*, whereas the remaining 1.1 million (9.61%) sites had the *A. majus* minor allele fixed in *A. molle*. For all such sites, the fixed *A. molle* allele was assigned ancestral, while the other *A. majus* allele derived.

**Pattern B:** *2,537,001 sites (21.99%) resolved.* Sites where both *A. majus* alleles are present in *A. molle*, and both species share the same major allele, i.e., allele frequency ≥ 0.5. For such sites, the major allele was assigned ancestral, since the minor allele in either taxa would most likely be derived due to new mutations being relatively rare.

**Pattern C:** *484,585 (4.21%) resolved.* Sites where both species are polymorphic, but only share 1 allele. In this case, the shared allele is considered ancestral, while the unique allele is considered derived.

The remaining 16.36% of bi-allelic sites in *A. majus* had no logical basis for assigning ancestral or derived alleles based on the information from *A. molle* (Patterns D–F in Table S4). The most unresolvable site-pattern (Pattern D: 1,418,581 or 12.30% of all bi-allelic sites), involved both species sharing the same 2 alleles, but the minor allele in one species was the major allele in the other. For 437,801 sites (3.80%), there was simply no sequence information available for *A. molle* (Pattern E in Table S4). Finally for the remaining 30,666 sites (0.26%), neither of the *A. majus* alleles was present in *A. molle* (Pattern F in Table S4), suggesting both alleles being derived in an unknown order in the ancestor to *A. majus*. For all the 1,887,048 unresolved bi-allelic sites (16.36%), we assumed the minor allele (< 0.5 allele frequency) in *A. majus* to be the most likely derived allele.

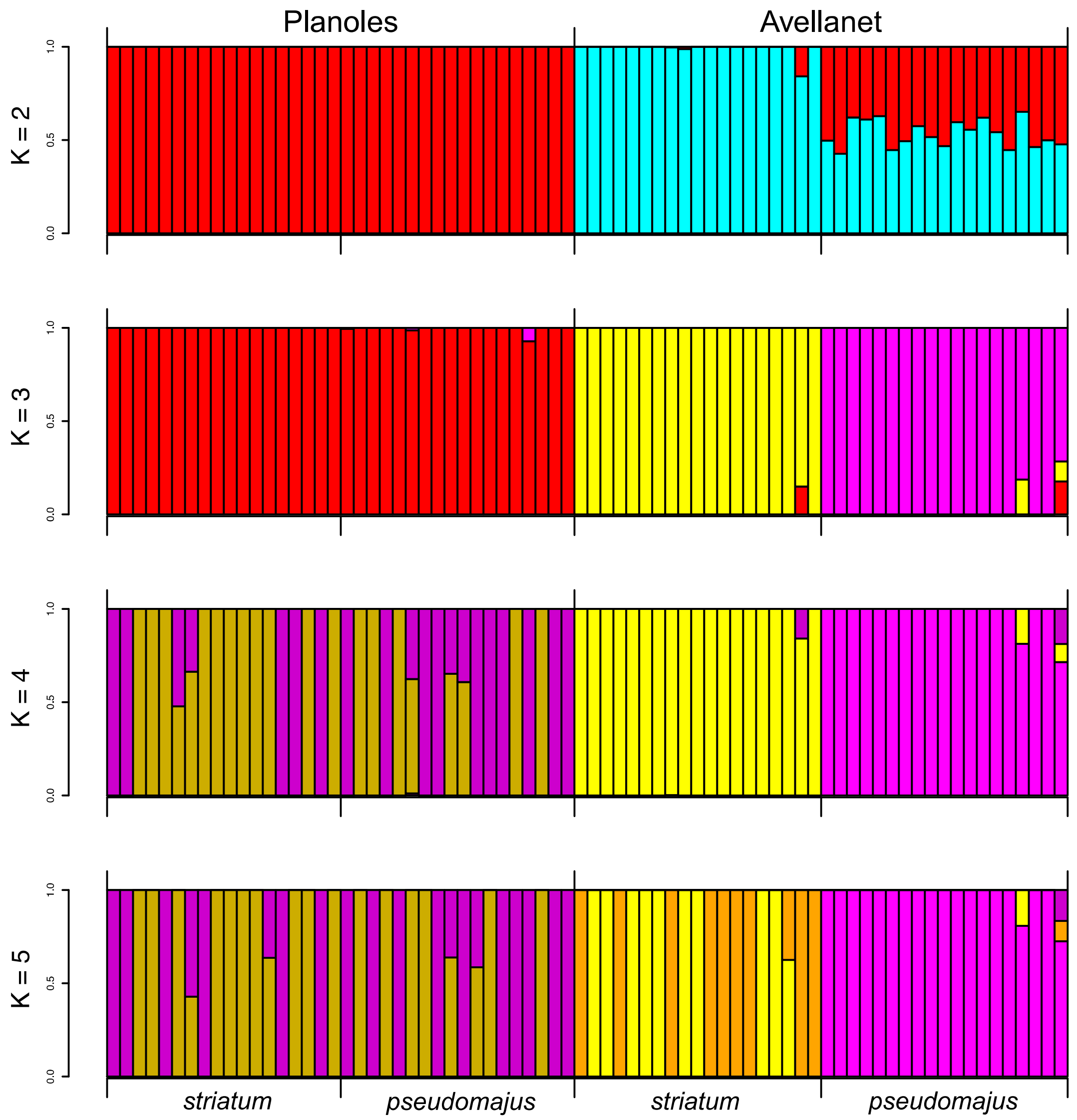


**Figure S1.** Admixture analysis of all 74 samples on 1,710,010 independent SNPs with *K* ranging from 2 to 5. *K*=3 describes the clearest ancestral grouping pattern, separating Avellanet into the 2 varieties, but grouping all Planoles samples together. None of the *K* values uniquely separate the var. *pseudomajus* and var. *striatum* samples at Planoles.

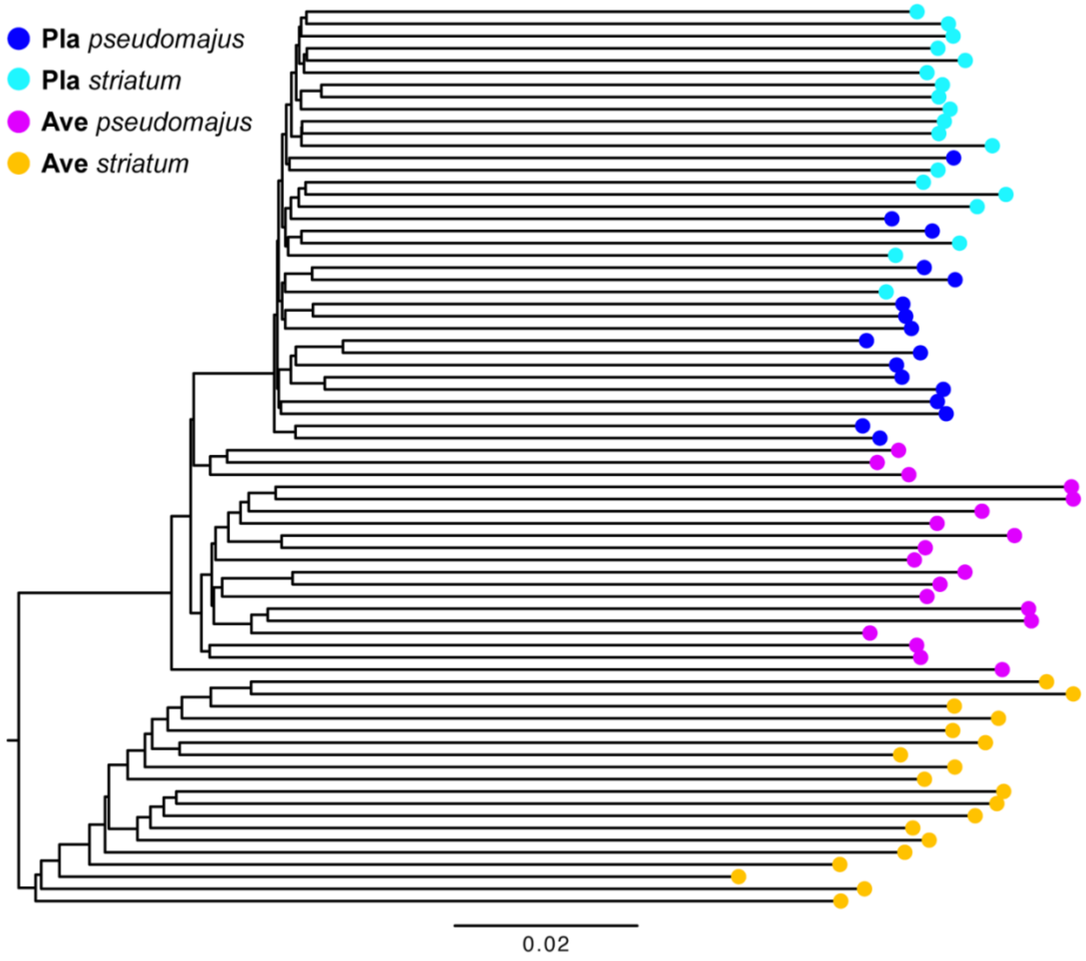


**Figure S2.** Neighbour-Joining tree inferred from 1,710,010 independent SNPs, which is used to calculate topology weighting for the whole genome.

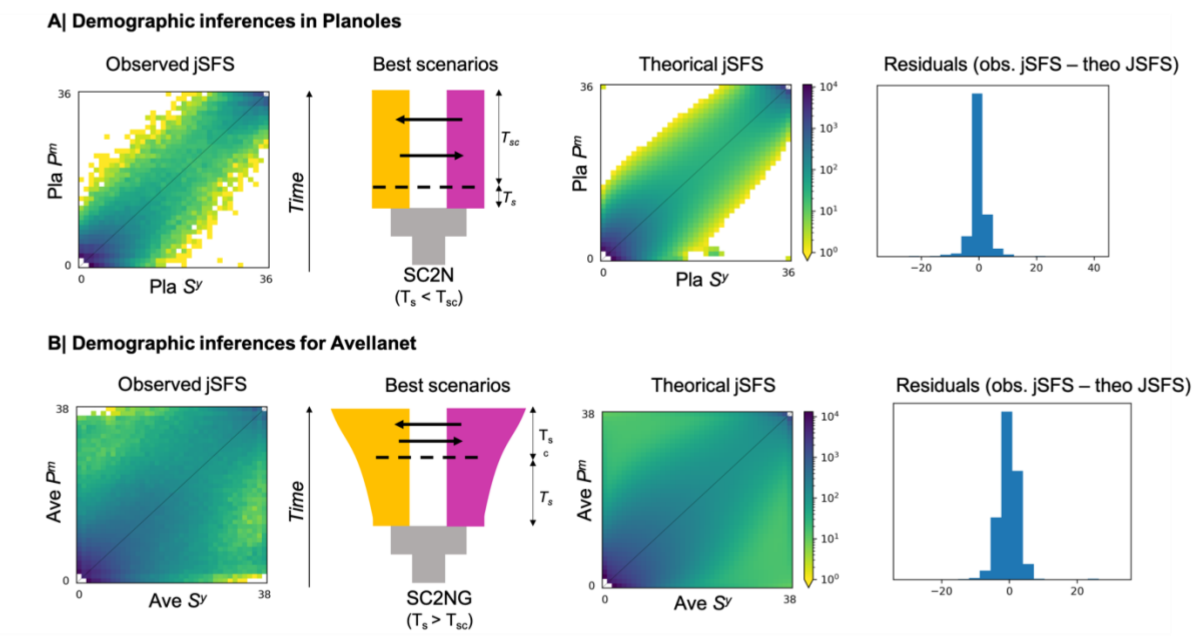


**Figure S3.** Results of the best demographic scenario inferred from *δaδi* analyses in **(A)** Planoles and **(B)** Avellanet. From left to right: the observed joint Site Frequency Spectrum ($jSFS_{obs}$), a schematic representation of the best-fitted demographic scenario, the theoretical spectrum ($jSFS_{theo}$) inferred from the best demographic scenario, and the distribution of residuals ($jSFS_{obs} - jSFS_{theo}$). For Planoles, the best-fitted scenario is a Secondary Contact with heterogeneous effective population size across the genome (SC2N). For Avellanet, the best-fitted scenario is a Secondary Contact with heterogeneous effective population size across the genome and exponential growth of the lineages (SC2NG).

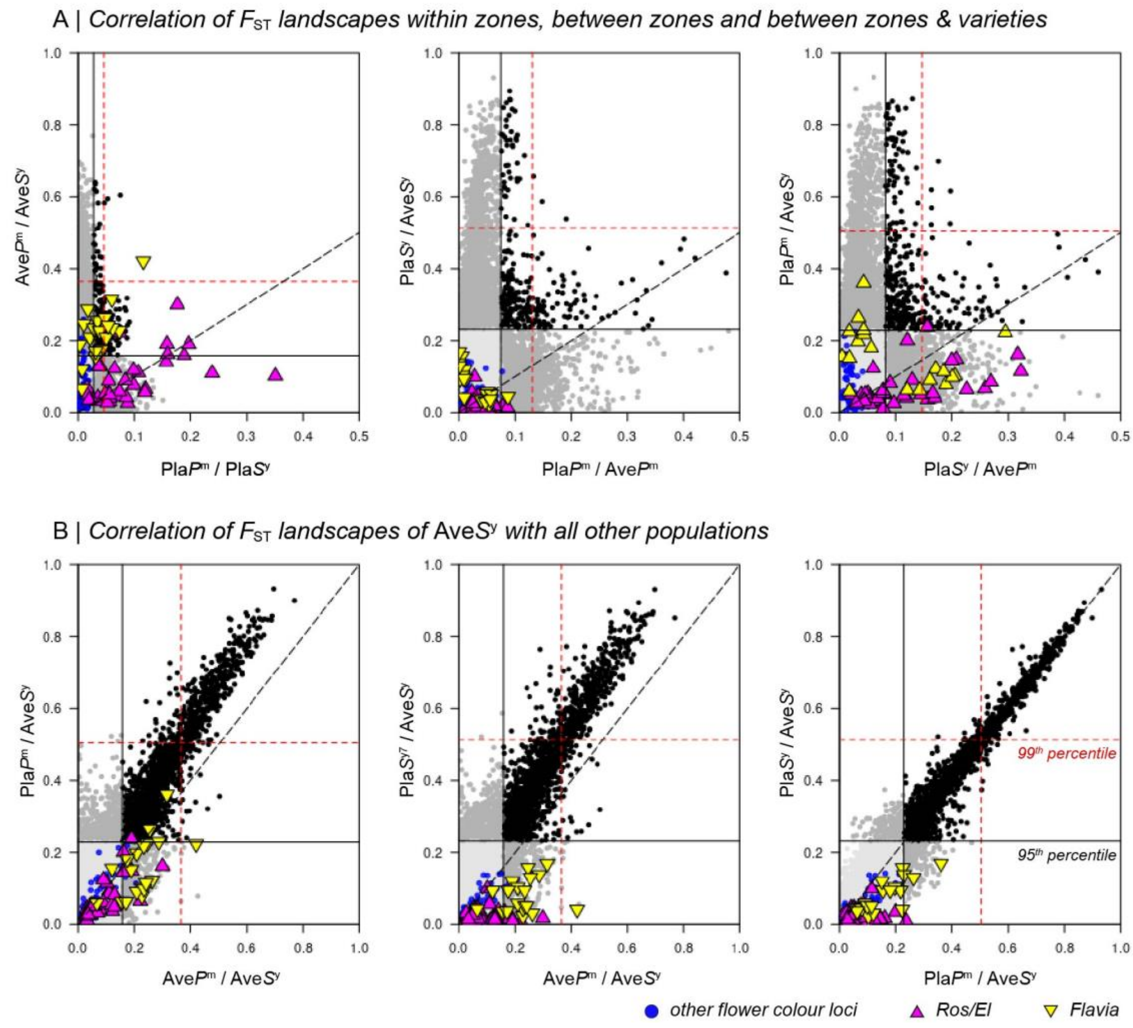


**Figure S4.** Correlations of pairwise Weir & Cockerham $F_{ST}$ in 10 Kbp non-overlapping windows. **(A)** The landscapes chosen for pairwise comparisons follow the same categories from top to bottom in Fig. 2 (Main Text) – i.e., within zones (Pla$P^m$/Pla$S^y$ vs. Ave$P^m$/Ave$S^y$), between zones (Pla$P^m$/Ave$P^m$ vs. Pla$S^y$/Ave$S^y$) and between zones & varieties (Pla$S^y$/Ave$P^m$ vs. Pla$P^m$/Ave$S^y$). **(B)** Correlations of $F_{ST}$ landscapes between Ave$S^y$ and all the other populations (Ave$P^m$/Ave$S^y$, Pla$P^m$/Ave$S^y$, Pla$S^y$/Ave$S^y$). 95$^{th}$ and 99$^{th}$ percentile thresholds are drawn in black and red within each plot, and black dots represent windows that are outliers (using 99$^{th}$ percentile) in both $F_{ST}$ landscapes. Magenta, yellow and blue dots represent windows that carry *Flavia*, *Ros/El* and other loci associated with flower colour. Pla: Planoles, Ave: Avellanet, $P^m$: magenta-coloured var. *pseudomajus*, $S^y$: yellow-coloured var. *striatum*.

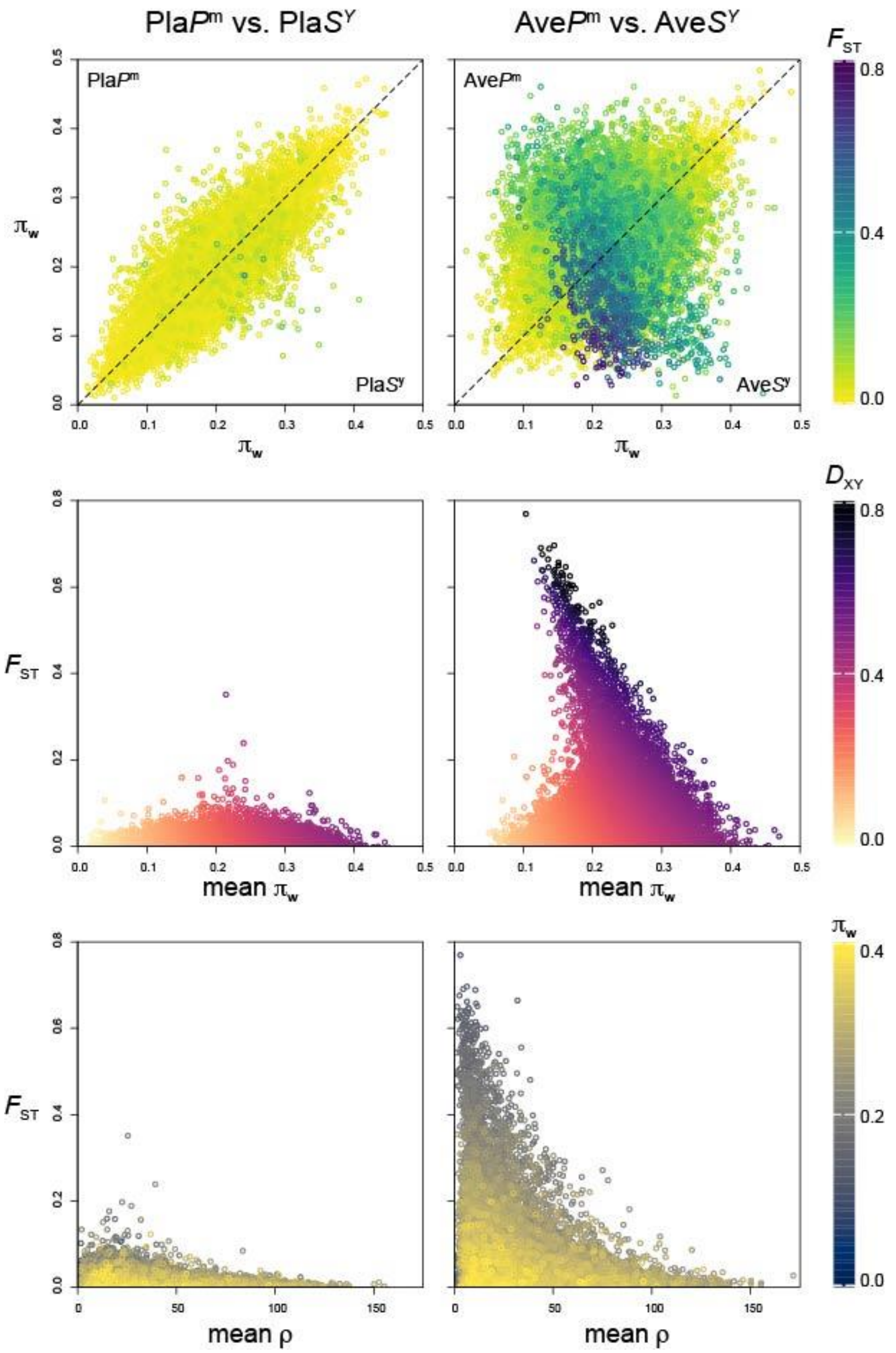


**Figure S5:** Relationships between population genetic estimates in 10 Kbp non-overlapping windows between both varieties at each hybrid zone. Top panel: $\pi_{\mathrm{w}}$ between the var. *pseudomajus* and var. *striatum*; coloured by $F_{\mathrm{ST}}$. Middle panel: $F_{\mathrm{ST}}$ vs. mean $\pi_{\mathrm{w}}$ ; coloured by $D_{\mathrm{XY}}$. Bottom panel: $F_{\mathrm{ST}}$ vs. mean $\rho$ (population recombination rate); coloured by mean $\pi_{\mathrm{w}}$. Pla: Planoles, Ave: Avellanet, $P^{\mathrm{m}}$: magenta-coloured var. *pseudomajus*, $S^{\mathrm{y}}$: yellow-coloured var. *striatum*.

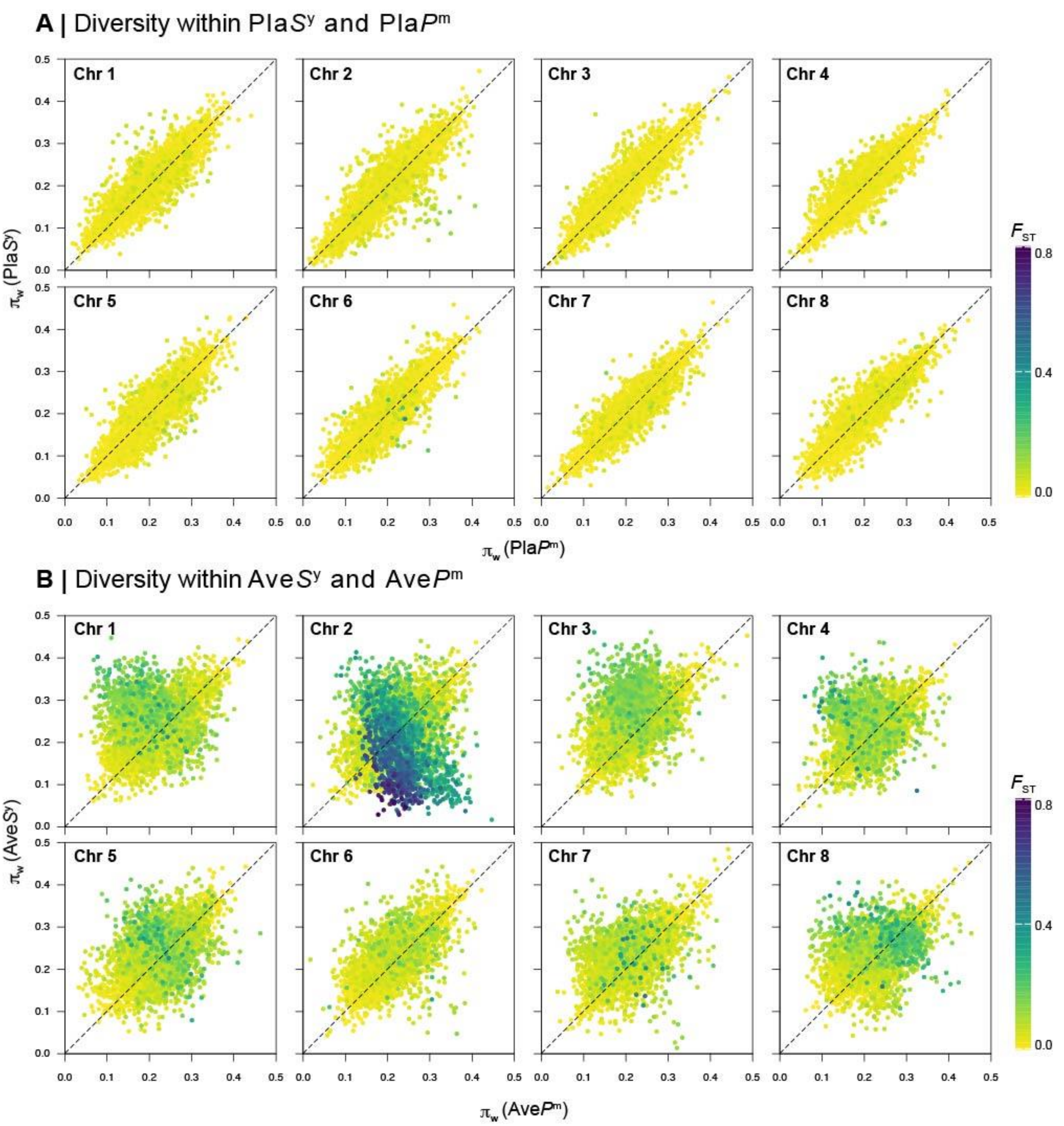


**Figure S6.** Relationship of diversity ($\pi_{\mathrm{w}}$) within var. *pseudomajus* and var. *striatum* in 10 Kbp non-overlapping windows at **(A)** Planoles and **(B)** Avellanet, shown for each chromosome. Dots are coloured by $F_{\mathrm{ST}}$. Pla: Planoles, Ave: Avellanet, $P^{\mathrm{m}}$: magenta-coloured var. pseudomajus, $S^{\mathrm{y}}$: yellow-coloured var. *striatum*.

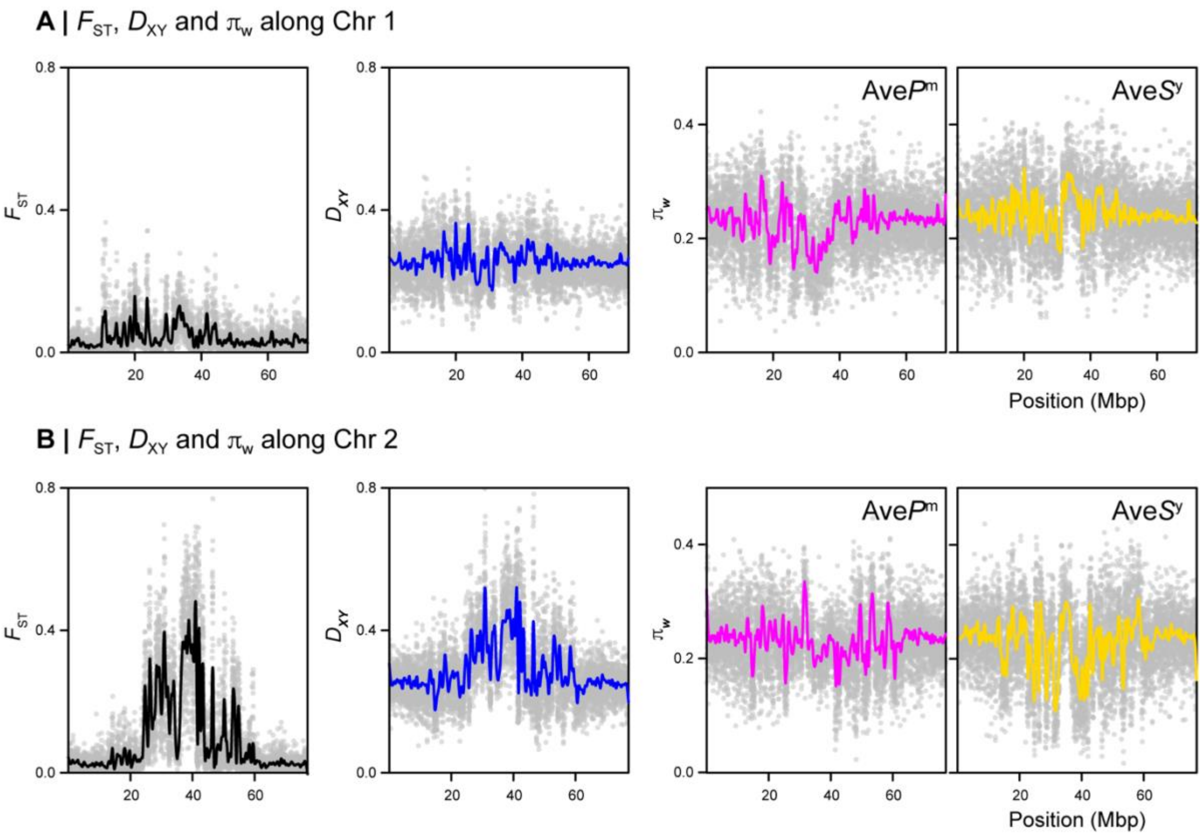


**Figure S7.** Population genetic parameters $F_{ST}$, $D_{XY}$ and $\pi_w$ along **(A)** Chr 1 and **(B)** Chr 2 for both varieties at Avellanet. Grey dots show estimates in 10 Kbp non-overlapping windows, while solid coloured lines show loess smoothed (span = 50 Kbp) estimates. Ave: Avellanet, $P^m$: magenta-coloured var. *pseudomajus*, $S^y$: yellow-coloured var. *striatum*.

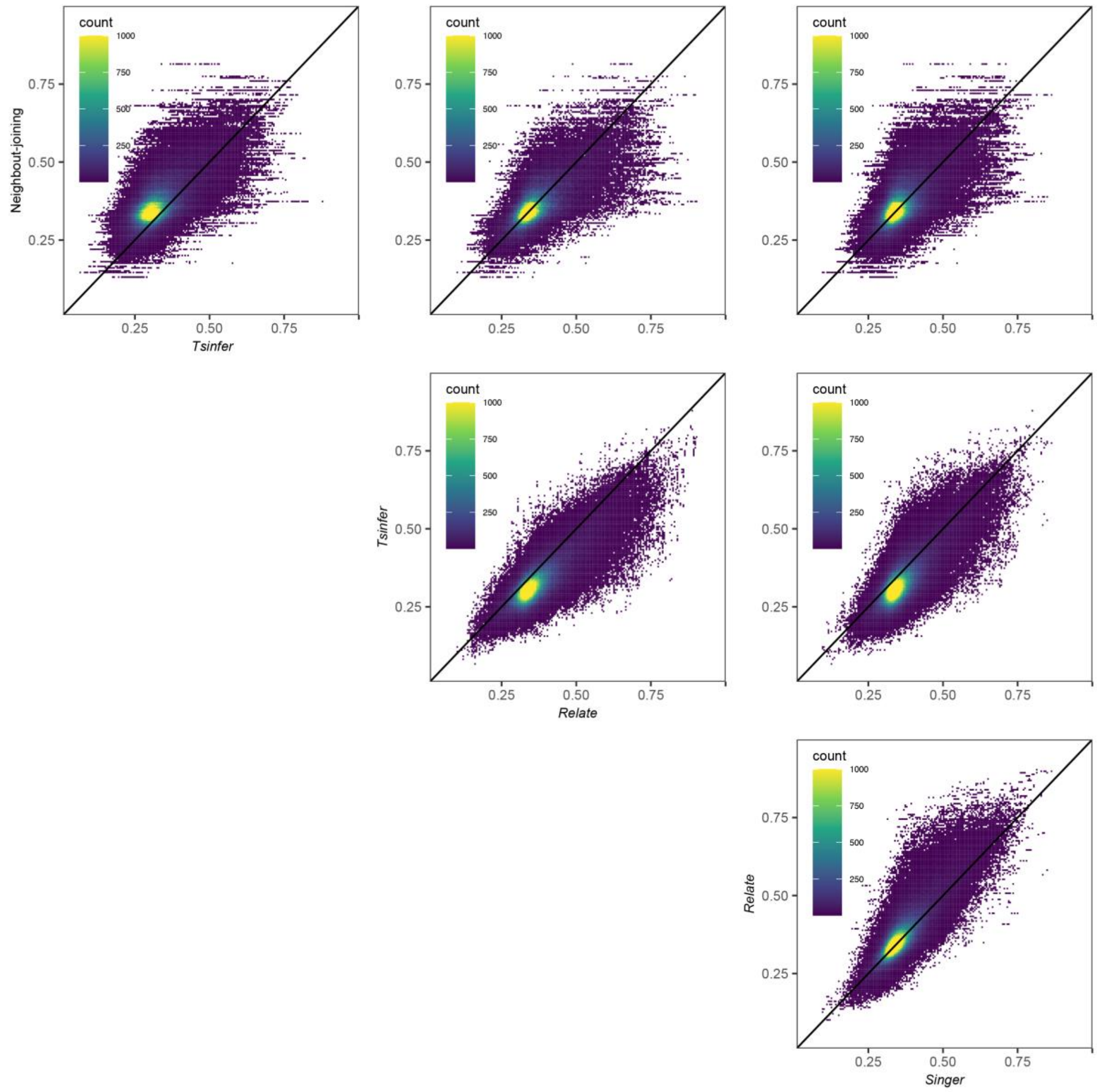


**Figure S8.** Correlation of $T_{geo}$ (clustered by geography) topology weights of trees inferred by different tree inference methods. Each square tile represents the number of trees in it.

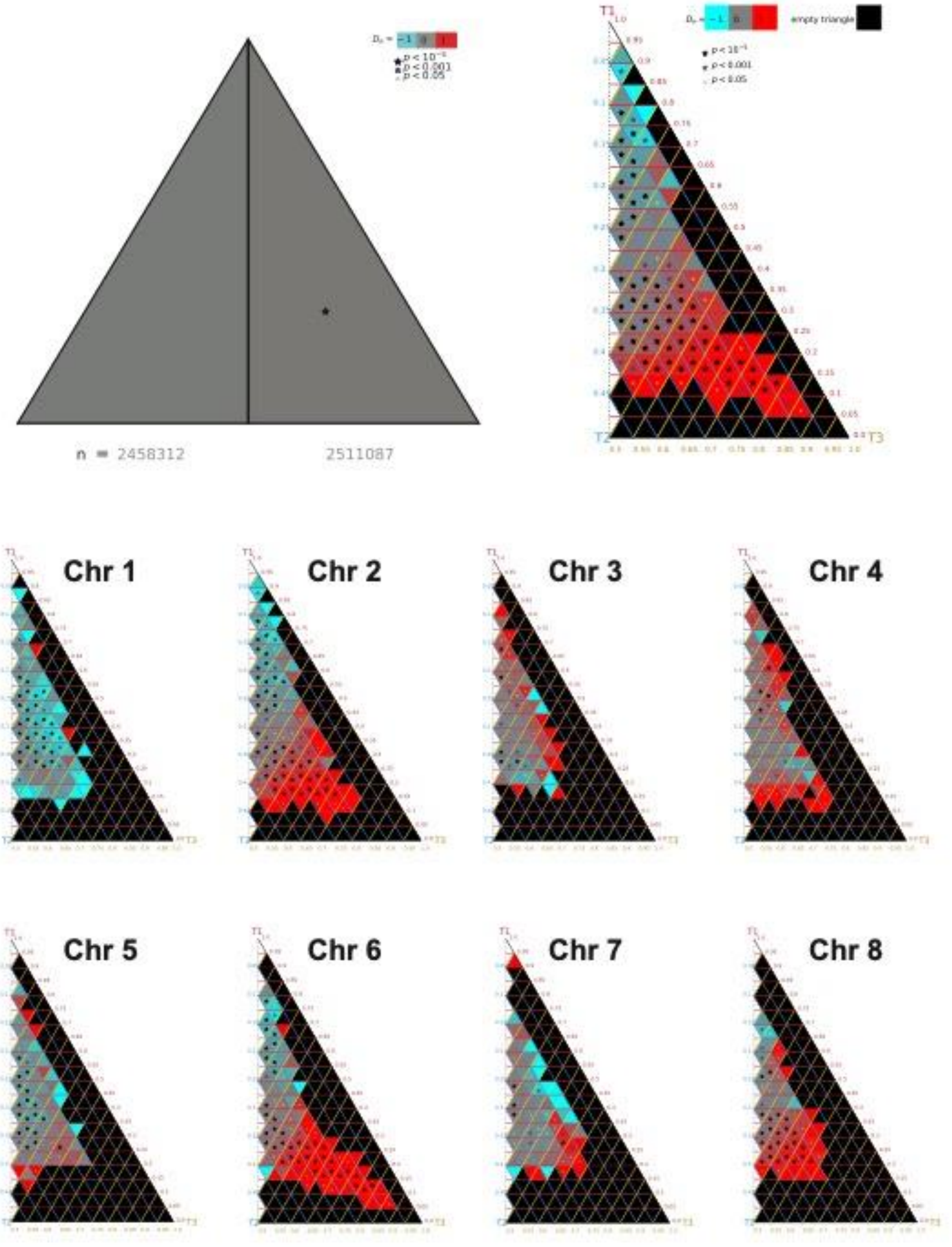


**Figure S9.** Left-right asymmetry from *TwisstNTern* analysis. Counts of trees in the left and right half-triangles, with asymmetry quantified using $D_{LR}$. Asterisks indicate significant asymmetry between corresponding left- and right-sided sub-triangles, suggesting excess of $T_{var}$ or $T_{alt}$ topologies.

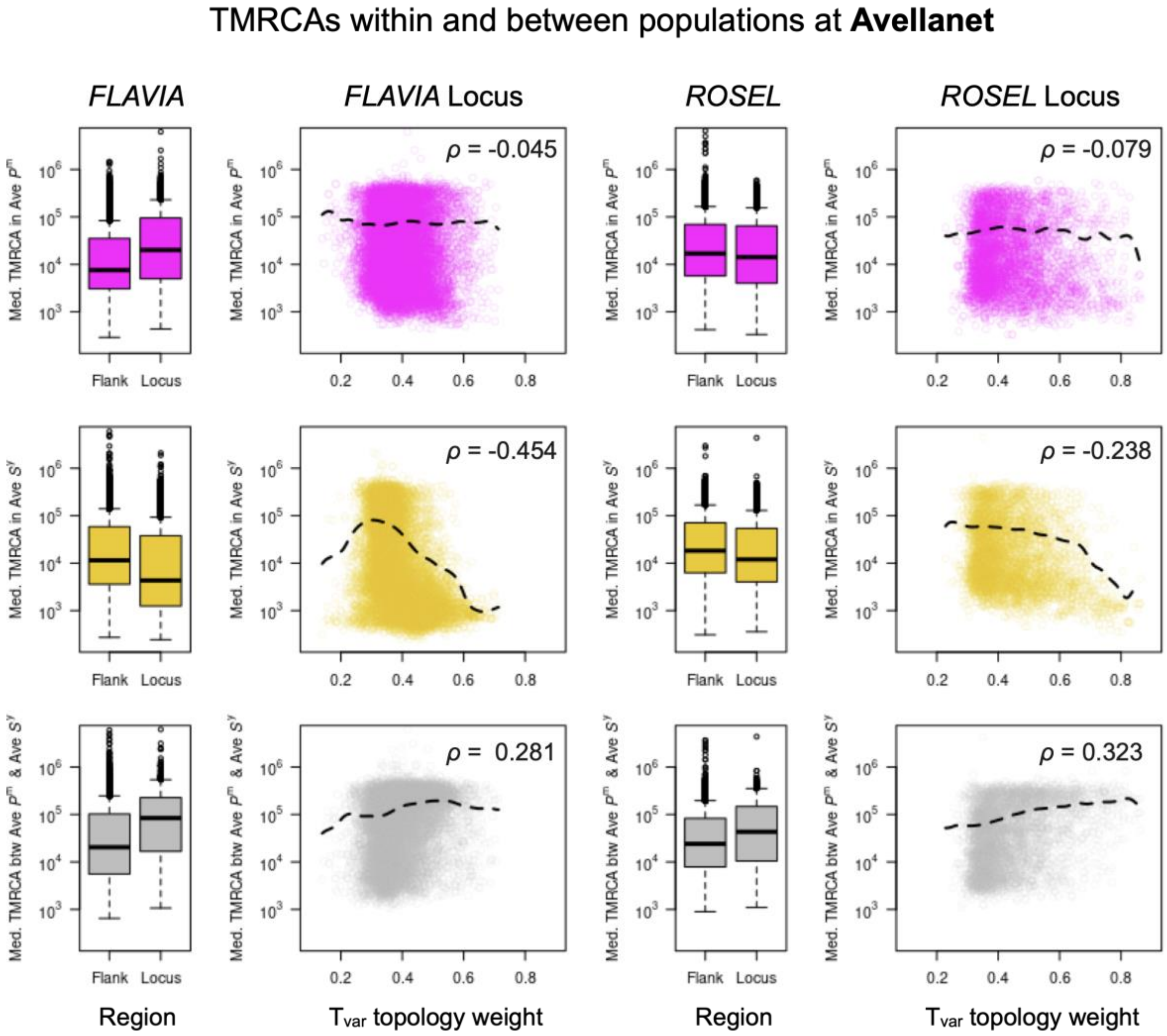


**Figure S10.** Coalescence times at *FLAVIA* and *ROS/EL* at the Avellanet populations. Boxplots for *FLAVIA* and *ROS/EL* show the median time to most recent common ancestor (TMRCA) within var. *pseudomajus* (top row), within var. *striatum* (middle row), and between var. *pseudomajus* and var. *striatum* (bottom row), all on a log scale. The left boxes ('Flank') show the TMRCAs in the flanking regions around the locus, while the right boxes ('Locus') show the values inside the locus. The scatterplots and dashed black lines show the full distribution and the overall smoothed trend between the $T_{var}$ weight and median TMRCA within the locus, (row are as indicated for the boxplots). $\rho$ is the correlation coefficient from a Spearman's rank correlation.

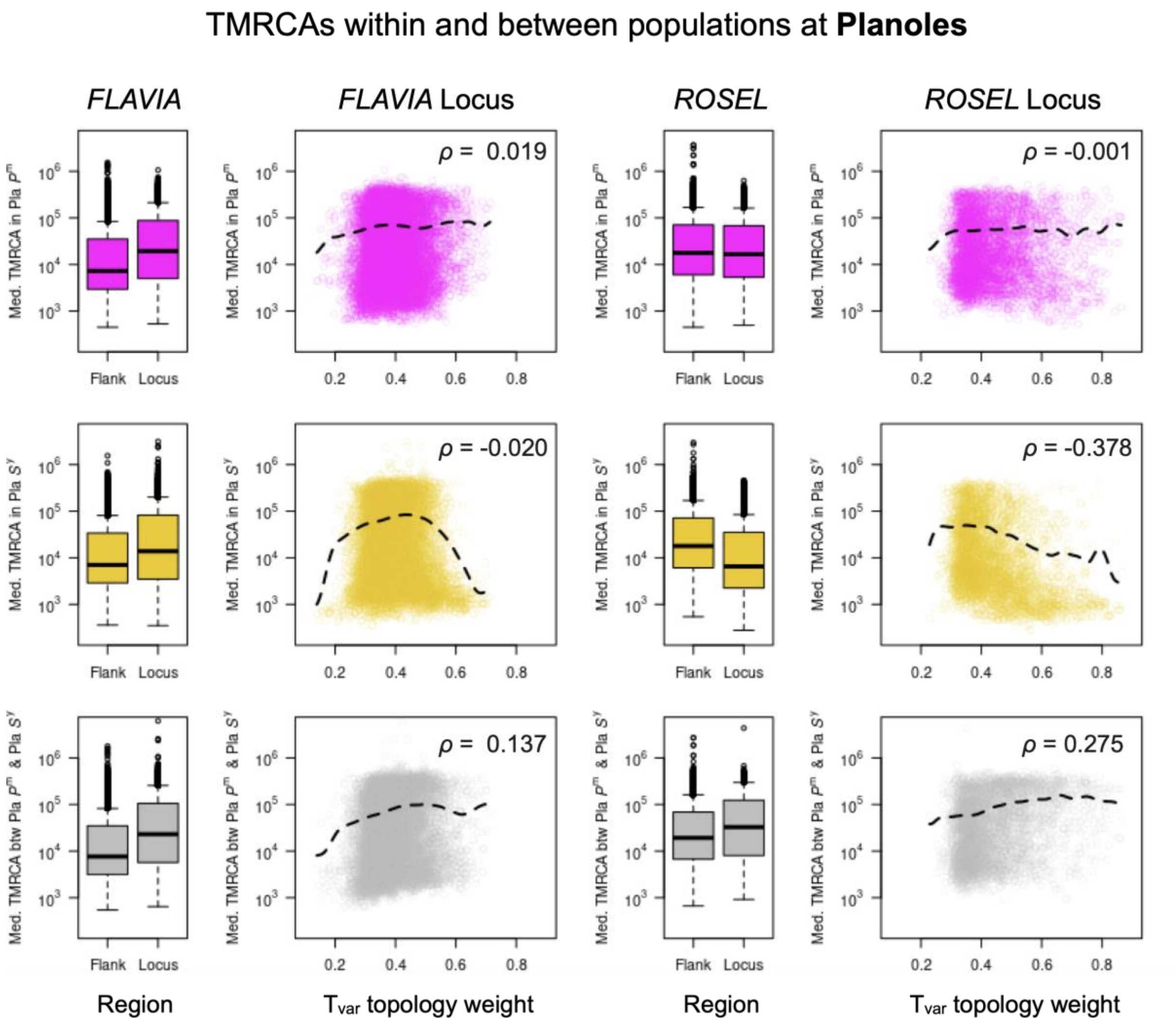


**Figure S11.** Coalescence times at *FLAVIA* and *ROS/EL* at the Planoles populations. Boxplots for *FLAVIA* and *ROS/EL* show the median time to most recent common ancestor (TMRCA) within var. *pseudomajus* (top row), within var. *striatum* (middle row), and between var. *pseudomajus* and var. *striatum* (bottom row), all on a log scale. The left boxes ('Flank') show the TMRCAs in the flanking regions around the locus, while the right boxes ('Locus') show the values inside the locus. The scatterplots and dashed black lines show the full distribution and the overall smoothed trend between the $T_{var}$ weight and median TMRCA within the locus, (row are as indicated for the boxplots). $\rho$ is the correlation coefficient from a Spearman's rank correlation.

**Table S1**. Locations and sequencing parameters for all samples. Pla: Planoles, Ave: Avellanet, *P*: var. *pseudomajus*, *S*: var. *striatum*.

| Plant ID | Population | | Latitude | Longitude | Altitude | Total reads | # Reads with correct barcodes | # Total reads mapped | Coverage | | Mapping Quality |
|---|---|---|---|---|---|---|---|---|---|---|---|
| | HZ | var. | | | | | | | mean | sd | |
| x3318 | Ave | *P* | 42.3265041 | 1.34596441 | 1180.397 | 3,188,122 | 3,129,590 | 3,105,569 | 0.79 | 5.30 | 43.86 |
| x3327 | Ave | *P* | 42.3253905 | 1.34795383 | 1176.437 | 2,371,661 | 2,329,207 | 2,297,793 | 0.58 | 3.97 | 43.79 |
| x3333 | Ave | *P* | 42.3254111 | 1.34793968 | 1175.417 | 3,981,670 | 3,922,672 | 3,870,878 | 0.98 | 5.78 | 43.51 |
| x3394 | Ave | *P* | 42.3265476 | 1.34605381 | 1182.238 | 5,286,233 | 5,198,347 | 5,108,647 | 1.30 | 7.35 | 43.64 |
| x4101 | Ave | *P* | 42.3254284 | 1.3479035 | 1173.368 | 4,997,162 | 4,898,390 | 4,835,944 | 1.23 | 6.37 | 43.47 |
| x4102 | Ave | *P* | 42.3253405 | 1.34797537 | 1174.698 | 4,002,073 | 3,935,551 | 3,892,352 | 0.99 | 7.08 | 43.11 |
| x4116 | Ave | *P* | 42.3253172 | 1.34792778 | 1173.042 | 3,873,173 | 3,780,189 | 3,755,484 | 0.96 | 5.67 | 43.98 |
| x4128 | Ave | *S* | 42.352512 | 1.32733675 | 1306.395 | 3,742,683 | 3,686,947 | 3,620,227 | 0.92 | 4.62 | 43.80 |
| x4129 | Ave | *S* | 42.3516395 | 1.32255535 | 1294.688 | 4,307,949 | 4,244,685 | 4,203,622 | 1.07 | 6.30 | 43.27 |
| x4134 | Ave | *S* | 42.3562788 | 1.31035377 | 1305.424 | 5,914,001 | 5,813,443 | 5,752,068 | 1.46 | 8.94 | 43.06 |
| x4137 | Ave | *S* | 42.3562711 | 1.31059675 | 1309.896 | 4,863,149 | 4,745,309 | 4,699,409 | 1.20 | 6.04 | 43.67 |
| x4140 | Ave | *S* | 42.3518215 | 1.32795229 | 1306.514 | 3,891,753 | 3,818,083 | 3,736,714 | 0.95 | 5.81 | 43.23 |
| x4148 | Ave | *S* | 42.3498082 | 1.32983188 | 1298.141 | 4,721,946 | 4,635,050 | 4,590,030 | 1.17 | 5.58 | 43.69 |
| x4149 | Ave | *S* | 42.3529544 | 1.31888622 | 1297.671 | 5,435,863 | 5,313,479 | 5,264,802 | 1.34 | 5.99 | 43.10 |
| x4150 | Ave | *S* | 42.3563588 | 1.31021196 | 1310.725 | 3,702,921 | 3,636,255 | 3,613,515 | 0.92 | 6.29 | 44.29 |
| x4153 | Ave | *S* | 42.3563674 | 1.31020257 | 1309.695 | 3,892,121 | 3,831,083 | 3,789,206 | 0.96 | 6.91 | 44.06 |
| x4155 | Ave | *S* | 42.3563344 | 1.31034666 | 1310.153 | 3,519,212 | 3,429,232 | 3,409,840 | 0.87 | 4.86 | 43.96 |
| x4158 | Ave | *S* | 42.3517021 | 1.32282928 | 1286.822 | 5,327,782 | 5,236,008 | 5,169,177 | 1.32 | 8.49 | 43.87 |
| x4159 | Ave | *S* | 42.3562196 | 1.3108351 | 1368.269 | 4,428,404 | 4,358,874 | 4,227,330 | 1.07 | 8.93 | 43.72 |
| x4161 | Ave | *P* | 42.3253486 | 1.34791663 | 1173.946 | 4,238,342 | 4,167,294 | 4,072,708 | 1.04 | 6.11 | 43.38 |
| x4178 | Ave | *P* | 42.3253335 | 1.34789948 | 1172.338 | 3,775,442 | 3,706,434 | 3,657,954 | 0.93 | 6.72 | 44.25 |
| x4359 | Ave | *P* | 42.3257335 | 1.3478534 | 1178.421 | 3,523,972 | 3,472,840 | 3,428,504 | 0.87 | 5.57 | 43.46 |
| x4392 | Ave | *S* | 42.3527129 | 1.31938189 | 1292.184 | 3,888,237 | 3,815,603 | 3,780,500 | 0.96 | 5.38 | 42.93 |

| | | | | | | | | | | | |
|---|---|---|---|---|---|---|---|---|---|---|---|
| x4429 | Ave | *P* | 42.3262125 | 1.34754946 | 1182.735 | 3,754,580 | 3,698,112 | 3,652,600 | 0.93 | 4.50 | 43.14 |
| x4459 | Ave | *P* | 42.3260837 | 1.34722933 | 1171.97 | 5,096,342 | 5,017,740 | 4,968,912 | 1.27 | 7.98 | 44.11 |
| x4463 | Ave | *S* | 42.3563577 | 1.31069132 | 1308.321 | 4,729,151 | 4,654,861 | 4,623,876 | 1.18 | 8.56 | 44.20 |
| x4477 | Ave | *S* | 42.3525811 | 1.32008953 | 1290.641 | 4,187,543 | 4,109,349 | 4,077,855 | 1.04 | 8.96 | 43.86 |
| x4491 | Ave | *S* | 42.3570279 | 1.30991165 | 1334.262 | 5,544,770 | 5,457,620 | 5,274,002 | 1.34 | 12.48 | 43.93 |
| x4570 | Ave | *S* | 42.3515774 | 1.32177665 | 1294.526 | 3,463,415 | 3,409,673 | 3,371,245 | 0.86 | 5.21 | 42.92 |
| x4579 | Ave | *S* | 42.3561238 | 1.31298015 | 1302.808 | 2,932,332 | 2,876,876 | 2,790,603 | 0.71 | 4.41 | 42.79 |
| x4580 | Ave | *S* | 42.3561336 | 1.3130905 | 1303.809 | 2,794,530 | 2,752,510 | 2,710,713 | 0.69 | 4.72 | 43.09 |
| x4585 | Ave | *P* | 42.3255331 | 1.34790478 | 1178.988 | 4,137,568 | 4,073,764 | 3,986,251 | 1.02 | 5.20 | 42.94 |
| x4593 | Ave | *P* | 42.3255351 | 1.34790367 | 1179.321 | 3,624,245 | 3,564,875 | 3,371,160 | 0.84 | 8.32 | 43.91 |
| x4608 | Ave | *P* | 42.3265208 | 1.34601267 | 1181.009 | 3,447,603 | 3,353,019 | 3,325,831 | 0.84 | 6.45 | 43.72 |
| x4624 | Ave | *P* | 42.326054 | 1.34714137 | 1171.926 | 5,276,887 | 5,170,031 | 5,105,650 | 1.29 | 10.77 | 43.93 |
| x4786 | Ave | *P* | 42.3263883 | 1.34558351 | 1183.084 | 3,124,533 | 3,072,873 | 3,025,727 | 0.77 | 5.37 | 44.02 |
| x4837 | Ave | *P* | 42.3250915 | 1.34802625 | 1176.246 | 3,006,238 | 2,932,758 | 2,857,202 | 0.72 | 5.89 | 43.77 |
| x4867 | Ave | *P* | 42.3263835 | 1.34565351 | 1181.582 | 2,081,727 | 2,047,913 | 2,009,962 | 0.51 | 2.38 | 42.61 |
| z0169 | Pla | *S* | 42.32448609 | 2.06488582 | 1225.909 | 3,331,472 | 3,257,556 | 3,136,601 | 0.80 | 6.43 | 44.77 |
| z0347 | Pla | *S* | 42.32502574 | 2.062415859 | 1230.095 | 2,600,606 | 2,562,974 | 2,477,371 | 0.63 | 4.24 | 44.03 |
| z0704 | Pla | *P* | 42.3236173 | 2.08200593 | 1184.011 | 2,265,609 | 2,226,435 | 2,198,175 | 0.56 | 3.02 | 43.52 |
| z0713 | Pla | *P* | 42.32365681 | 2.081934964 | 1182.894 | 3,402,360 | 3,336,764 | 3,302,472 | 0.84 | 5.82 | 43.32 |
| z0847 | Pla | *S* | 42.32463343 | 2.06416347 | 1229.266 | 2,662,730 | 2,620,318 | 2,586,796 | 0.66 | 5.02 | 45.00 |
| z0852 | Pla | *S* | 42.32446099 | 2.065055654 | 1222.145 | 3,522,296 | 3,460,620 | 3,350,803 | 0.85 | 7.37 | 44.35 |
| z0854 | Pla | *S* | 42.3246058 | 2.06428122 | 1229.051 | 3,398,868 | 3,343,220 | 3,227,563 | 0.82 | 6.50 | 44.71 |
| z1778 | Pla | *P* | 42.32322484 | 2.082552166 | 1177.311 | 4,023,628 | 3,890,344 | 3,853,827 | 0.98 | 8.99 | 44.83 |
| z1802 | Pla | *S* | 42.32451883 | 2.064910802 | 1224.598 | 3,804,550 | 3,745,410 | 3,673,814 | 0.94 | 7.19 | 45.03 |
| z2008 | Pla | *S* | 42.3250673 | 2.062047337 | 1238.035 | 3,300,071 | 3,244,531 | 3,148,951 | 0.80 | 6.35 | 44.80 |
| z2018 | Pla | *S* | 42.32451109 | 2.064902788 | 1225.374 | 4,536,347 | 4,459,591 | 4,328,480 | 1.10 | 7.73 | 44.36 |
| z2020 | Pla | *S* | 42.32503701 | 2.06272073 | 1237.244 | 3,693,881 | 3,626,029 | 3,442,237 | 0.87 | 5.91 | 43.75 |

| | | | | | | | | | | | |
|---|---|---|---|---|---|---|---|---|---|---|---|
| z2046 | Pla | *P* | 42.32322195 | 2.082707483 | 1195.883 | 2,856,165 | 2,786,839 | 2,738,491 | 0.70 | 4.79 | 45.05 |
| z2281 | Pla | *P* | 42.32368318 | 2.081933918 | 1159.164 | 2,278,684 | 2,235,650 | 2,167,369 | 0.55 | 2.94 | 44.38 |
| z2283 | Pla | *P* | 42.3235961 | 2.081999912 | 1182.457 | 2,753,499 | 2,714,511 | 2,639,088 | 0.67 | 3.12 | 43.28 |
| z2286 | Pla | *P* | 42.323607 | 2.081954368 | 1185.514 | 2,369,109 | 2,304,211 | 2,208,590 | 0.56 | 2.82 | 43.27 |
| z2287 | Pla | *P* | 42.32360853 | 2.081924361 | 1180.977 | 2,358,238 | 2,304,338 | 2,202,698 | 0.56 | 2.84 | 43.82 |
| z2290 | Pla | *P* | 42.32361059 | 2.081978279 | 1184.116 | 3,135,858 | 3,071,886 | 2,985,428 | 0.76 | 4.75 | 44.93 |
| z2292 | Pla | *P* | 42.32359454 | 2.081963851 | 1184.393 | 3,248,824 | 3,182,852 | 3,054,138 | 0.77 | 4.88 | 44.92 |
| z2293 | Pla | *P* | 42.32360997 | 2.081978164 | 1182.295 | 2,129,244 | 2,092,966 | 2,043,766 | 0.52 | 3.51 | 44.68 |
| z2294 | Pla | *P* | 42.32359948 | 2.081961153 | 1183.949 | 3,245,003 | 3,177,589 | 3,085,916 | 0.78 | 4.67 | 44.18 |
| z2295 | Pla | *P* | 42.32360819 | 2.081973936 | 1183.365 | 3,343,971 | 3,293,189 | 3,207,905 | 0.81 | 4.69 | 44.42 |
| z2296 | Pla | *P* | 42.32360588 | 2.081925299 | 1181.766 | 2,350,694 | 2,297,670 | 2,202,817 | 0.56 | 5.46 | 44.12 |
| z2298 | Pla | *P* | 42.32361395 | 2.081979136 | 1181.428 | 3,902,911 | 3,840,117 | 3,767,598 | 0.96 | 8.04 | 44.67 |
| z2300 | Pla | *P* | 42.32360116 | 2.081929359 | 1183.732 | 3,175,981 | 3,105,901 | 2,997,729 | 0.76 | 3.42 | 42.94 |
| z2706 | Pla | *S* | 42.32458183 | 2.064503601 | 1233.894 | 3,240,057 | 3,185,969 | 3,120,068 | 0.80 | 3.25 | 44.17 |
| z2711 | Pla | *S* | 42.32504359 | 2.061967047 | 1235.348 | 5,045,346 | 4,964,032 | 4,867,405 | 1.24 | 8.07 | 44.45 |
| z3891 | Pla | *S* | 42.32512037 | 2.062697473 | 1467.332 | 3,859,686 | 3,798,512 | 3,748,425 | 0.95 | 5.75 | 44.61 |
| z4710 | Pla | *P* | 42.32364551 | 2.081975459 | 1177.4 | 2,361,709 | 2,322,695 | 2,203,640 | 0.56 | 3.64 | 44.18 |
| z4712 | Pla | *P* | 42.32364214 | 2.081965884 | 1176.806 | 2,802,140 | 2,749,078 | 2,609,075 | 0.66 | 4.00 | 44.29 |
| z4758 | Pla | *S* | 42.3248285 | 2.063343366 | 1300.989 | 5,301,316 | 5,189,766 | 5,132,422 | 1.31 | 10.83 | 44.73 |
| z4783 | Pla | *S* | 42.32459417 | 2.064393839 | 1228.686 | 3,220,924 | 3,165,208 | 3,123,531 | 0.80 | 5.27 | 44.34 |
| z4793 | Pla | *S* | 42.32491715 | 2.063297364 | 1222.045 | 2,257,174 | 2,220,722 | 2,178,325 | 0.55 | 4.28 | 44.72 |
| z4797 | Pla | *S* | 42.32166099 | 2.070664005 | 1203.821 | 2,568,966 | 2,487,622 | 2,447,585 | 0.62 | 4.29 | 44.80 |
| z5444 | Pla | *S* | 42.32488623 | 2.063280532 | 1220.282 | 3,209,147 | 3,152,037 | 3,128,019 | 0.80 | 5.27 | 44.79 |
| z5451 | Pla | *S* | 42.3245906 | 2.06439466 | 1213.522 | 3,030,447 | 2,949,741 | 2,916,628 | 0.74 | 3.73 | 44.63 |

**Table S2:** Results of the demographic modelling for the five best replicate (lower AIC value) of each the eight tested scenarios. The table shows, in order of appearance, the focal studied population, the model tested, its AIC, the population mutation rate of the ancestral population (Θ), and the following parameter estimated by the model that are scaled to Θ: the change in ancestral population size (Na), the population size of the first ($Ne_1 = S^y$) and second ($Ne_2 = P^m$) population after the split, the exponential population growth parameter in $S^y$ ($b_1$) and $P^m$ ($b_2$), the Hill-Robertson effect (hrf) corresponding to the fraction by which Ne is reduced in region strongly affected by selection at linked site, the time at which ancestral variation in effective size occurred ($T_a$), the time at which derived population have split ($T_s$), the time of secondary contact ($T_{sc}$), and the fraction of the genome affected by reduced Ne (Q), and the fraction of derived mutation miss oriented in the unfolded spectrum (O). The best model is highlighted in bold.

| Population | Model | AIC | Theta | Na | $Ne_1$ | $Ne_2$ | b1 | b2 | hrf | $M_{1>2}$ | $M_{2>1}$ | $T_a$ | $T_s$ | $T_{sc}$ | Q | O | Ts/Tsc |
|---|---|---|---|---|---|---|---|---|---|---|---|---|---|---|---|---|---|
| Avellanet | **SCA2N** | **17221** | **8505** | **35.46** | **2.62** | **1.67** | **0.02** | **-** | **-** | **1.00** | **8.44** | **1.65** | **0.00** | **0.40** | **0.01** | **0.93** | **3365.85** |
| | SCA2NG | 17462 | 6920 | 33.96 | 3.92 | 1.68 | 0.83 | 1.48 | 0.16 | 0.84 | 5.75 | 2.19 | 0.03 | 0.59 | 0.01 | 0.93 | 18.08 |
| | SCAG | 18414 | 9715 | 17.06 | 6.74 | 2.72 | 0.09 | 0.28 | - | 7.09 | 11.90 | 1.11 | 0.33 | 0.19 | - | 0.93 | 0.56 |
| | SCA | 18995 | 6114 | 11.02 | 2.14 | 2.74 | - | - | - | 0.92 | 1.66 | 3.48 | 0.02 | 0.06 | - | 0.93 | 2.41 |
| | SIA2NG | 18396 | 7129 | 9.71 | 2.34 | 2.04 | 2.53 | 5.43 | 0.02 | - | - | 2.92 | 0.00 | - | 0.94 | 0.93 | - |
| | SIA2N | 24261 | 14828 | 4.60 | 8.18 | 3.64 | - | - | 0.83 | - | - | 0.09 | 0.18 | - | 0.26 | 0.95 | - |
| | SIAG | 18444 | 4826 | 14.04 | 0.16 | 0.96 | 11.14 | 0.46 | - | - | - | 4.75 | 0.01 | - | - | 0.92 | - |
| | SIA | 18418 | 6987 | 9.67 | 0.00 | 0.01 | - | - | - | - | - | 3.08 | 0.00 | - | - | 0.92 | - |
| Planoles | SCA2N | 26078 | 25952 | 15.00 | 0.96 | 1.39 | - | - | 0.20 | 4.71 | 3.05 | 4.76 | 0.28 | 0.26 | 0.37 | 0.97 | 0.93 |
| | **SCA2NG** | **26036** | **15638** | **15.16** | **1.35** | **2.07** | **1.53** | **0.61** | **0.15** | **2.33** | **2.67** | **8.55** | **0.41** | **0.25** | **0.18** | **0.96** | **0.60** |
| | SCA | 49246 | 48581 | 42.57 | 0.41 | 0.41 | 5.63 | 6.07 | | | | 1.43 | 0.01 | 0.29 | | 0.97 | 38.41 |
| | SCAG | 48740 | 10656 | 17.15 | 2.45 | 2.40 | 0.71 | 0.84 | 1.41 | 1.36 | | 13.77 | 0.65 | 0.44 | | 0.97 | 0.68 |
| | SIA2N | 49784 | 102652 | 0.40 | 0.07 | 0.07 | - | - | 0.01 | - | - | 0.00 | 0.11 | - | 0.19 | 1.00 | |
| | SIA2NG | 47075 | 104479 | 0.02 | 0.01 | 0.01 | 2.42 | 2.18 | 0.20 | - | - | 0.00 | 0.00 | - | 0.18 | 1.00 | |
| | SIA | 69406 | 104153 | 0.33 | 0.05 | 0.05 | | | | | | 0.01 | 0.00 | | | 1.00 | |
| | SIAG | 53104 | 195956 | 0.50 | 0.09 | 0.07 | 0.04 | 0.05 | | | | 6.44 | 0.00 | | | 1.00 | |

**Table S3.** Overview of number of sites at the end of step of variant filtering by *bcftools*, along with the representative codes to implement each step. The final 11,574,426 sites were used as candidate positions for imputation and inference of SNPs by *STITCH*.

<table>
<tr><th>Step</th><th>Dataset</th><th>Representative Code</th><th># sites</th></tr>
<tr><td>1</td><td>All sites</td><td>bcftools mpileup --annotate AD,ADF,ADR,DP,QS,SP -d 500 |<br>bcftools call --annotate GQ,GP -m</td><td>495,577,994</td></tr>
<tr><td>2</td><td>Variant sites within 5bp of INDELs removed and only bi-allelic SNPs retained.</td><td>bcftools view -m2 -M2 -v snps -e "AC==0 || AC==AN"</td><td>17,107,390</td></tr>
<tr><td>3</td><td>SNPs removed based on depth and quality.</td><td>bcftools filter -e "INFO/DP>130 | QUAL<20 | MQ<20"</td><td>12,727,484</td></tr>
<tr><td>4</td><td>SNPs with ≤ 0.8 missing fraction retained.</td><td>bcftools filter -e "F_MISSING>0.8"</td><td>11,574,426</td></tr>
</table>

**Table S4.** Overview of number of sites before and after calling and imputation of SNPs by *STITCH*. INFO score, computed by *STITCH*, reflects confidence in imputation accuracy. Generally, INFO ≥ 0.8 is considered high confidence (93.9% of all sites).

| Step | Types of sites | # sites (%) |
|---|---|---|
| **Before *STITCH* imputation** | Initial candidate variant sites (inferred from *bcftools*) | 11,574,426 – |
| **After *STITCH* imputation** | Invariant sites | 41,396 (0.4%) |
| | Variant sites (all) | 11,533,030 (99.6%) |
| | Variant sites (INFO ≥ 0.8) | 10,873,003 (93.94%) |
| | Variant sites (INFO ≥ 0.6) | 11,443,277 (98.9%) |
| | Variant sites (INFO ≥ 0.4) | 11,526,221 (99.6%) |

**Table S5.** Overview of number of sites where *A. majus* alleles could be polarised into ancestral or derived, based on alleles present in *Antirrhinum molle*. For unresolved sites, major allele in *A.majus* was considered ancestral. *unres.*: Unresolved

| Type | Site Description | Ancestral Allele | Derived Allele | No. of sites (%) for each type | Total no. of resolved/unresolved sites (%) | |
|---|---|---|---|---|---|---|
| **A** | One of the *A.majus* allele is fixed in *A molle*. | FIXED | Other | 6,624,396 (57.4%) | 9,645,982 (83.6%) | Resolved sites |
| **B** | Both *A.majus* alleles present in *A.molle* and they have the same major allele (frequency ≥ 0.5). | MAJOR | MINOR | 2,537,001 (22.0%) | | |
| **C** | Both *A.majus* and *A.molle* are polymorphic, but only share 1 allele. | SHARED | Other | 484,585 (4.2%) | | |
| **D** | Both *A.majus* alleles present in *A.molle*, but they do **not** have the same major allele (frequency ≥ 0.5). | *unres.* | *unres.* | 1,418,581 (12.3%) | 1,887,048 (16.4%) | Unresolved sites |
| **E** | No genotype information for *A.molle*. | *unres.* | *unres.* | 437,801 (3.8%) | | |
| **F** | No shared allele between A.majus and *A.molle*. | *unres.* | *unres.* | 30,666 (0.3%) | | |

**Table S6.** Correlation (Spearman's rho) between population genetic parameters in 10Kb non-overlapping windows. Values ≥ 0.60 are highlighted in green. Pla: Planoles, Ave: Avellanet, $P^{m}$: magenta-coloured var. *pseudomajus*, $S^{y}$: yellow-coloured var. *striatum*.

| $\pi_w$ | | | | $D_{XY}$ | | | | | | $F_{ST}$ | | | | | | | |
|---|---|---|---|---|---|---|---|---|---|---|---|---|---|---|---|---|---|
| Ave$P^{m}$ | Ave$S^{y}$ | Pla$P^{m}$ | Pla$S^{y}$ | Ave$P^{m}$ – Ave$S^{y}$ | Ave$P^{m}$ – Pla$P^{m}$ | Ave$P^{m}$ – Pla$S^{y}$ | Ave$S^{y}$ – Pla$P^{m}$ | Ave$S^{y}$ – Pla$S^{y}$ | Pla$P^{m}$ – Pla$S^{y}$ | Ave$P^{m}$ – Ave$S^{y}$ | Ave$P^{m}$ – Pla$P^{m}$ | Ave$P^{m}$ – Pla$S^{y}$ | Ave$S^{y}$ – Pla$P^{m}$ | Ave$S^{y}$ – Pla$S^{y}$ | Pla$P^{m}$ – Pla$S^{y}$ | | |
| 0.26 | 0.13 | 0.38 | 0.39 | 0.07 | 0.3 | 0.29 | 0.07 | 0.07 | 0.39 | -0.3 | -0.23 | -0.27 | -0.37 | -0.37 | -0.08 | **mean recomb. rate** | |
| | 0.47 | 0.66 | 0.64 | 0.73 | 0.89 | 0.89 | 0.63 | 0.63 | 0.67 | -0.07 | 0.02 | 0.04 | 0 | 0 | -0.01 | Ave$P^{m}$ | $\pi_w$ |
| | | 0.32 | 0.31 | 0.77 | 0.42 | 0.41 | 0.73 | 0.73 | 0.32 | 0.1 | -0.03 | -0.01 | 0.15 | 0.16 | -0.04 | Ave$S^{y}$ | $\pi_w$ |
| | | | 0.89 | 0.42 | 0.88 | 0.84 | 0.53 | 0.5 | 0.97 | -0.17 | -0.17 | -0.12 | -0.28 | -0.24 | -0.02 | Pla$P^{m}$ | $\pi_w$ |
| | | | | 0.39 | 0.82 | 0.87 | 0.47 | 0.52 | 0.97 | -0.19 | -0.13 | -0.17 | -0.26 | -0.29 | -0.01 | Pla$S^{y}$ | $\pi_w$ |
| | | | | | 0.62 | 0.61 | 0.92 | 0.91 | 0.42 | 0.36 | 0.07 | 0.11 | 0.35 | 0.35 | -0.02 | Ave$P^{m}$ – Ave$S^{y}$ | $D_{XY}$ |
| | | | | | | 0.97 | 0.64 | 0.62 | 0.87 | -0.11 | 0.08 | 0.08 | -0.1 | -0.09 | 0.01 | Ave$P^{m}$ – Pla$P^{m}$ | $D_{XY}$ |
| | | | | | | | 0.61 | 0.64 | 0.88 | -0.11 | 0.07 | 0.1 | -0.09 | -0.09 | 0.02 | Ave$P^{m}$ – Pla$S^{y}$ | $D_{XY}$ |
| | | | | | | | | 0.97 | 0.52 | 0.33 | 0.06 | 0.1 | 0.41 | 0.4 | -0.02 | Ave$S^{y}$ – Pla$P^{m}$ | $D_{XY}$ |
| | | | | | | | | | 0.52 | 0.32 | 0.06 | 0.11 | 0.39 | 0.41 | -0.01 | Ave$S^{y}$ – Pla$S^{y}$ | $D_{XY}$ |
| | | | | | | | | | | -0.18 | -0.15 | -0.13 | -0.27 | -0.26 | 0.05 | Pla$P^{m}$ – Pla$S^{y}$ | $D_{XY}$ |
| | | | | | | | | | | | 0.19 | 0.24 | 0.64 | 0.63 | 0.04 | Ave$P^{m}$ – Ave$S^{y}$ | $F_{ST}$ |
| | | | | | | | | | | | | 0.67 | 0.31 | 0.24 | 0.11 | Ave$P^{m}$ – Pla$P^{m}$ | $F_{ST}$ |
| | | | | | | | | | | | | | 0.29 | 0.38 | 0.16 | Ave$P^{m}$ – Pla$S^{y}$ | $F_{ST}$ |
| | | | | | | | | | | | | | | 0.86 | 0.07 | Ave$S^{y}$ – Pla$P^{m}$ | $F_{ST}$ |
| | | | | | | | | | | | | | | | 0.07 | Ave$S^{y}$ – Pla$S^{y}$ | $F_{ST}$ |
| | | | | | | | | | | | | | | | | Pla$P^{m}$ – Pla$S^{y}$ | $F_{ST}$ |

**Table S7.** Correlation (Spearman's rho) between topology weights inferred from different tree inference methods. Values ≥ 0.60 are highlighted in green.

| **Tg - Geography Topology** | | | | |
|---|---|---|---|---|
| *NJ* | *tsinfer* | *Relate* | *Singer* | |
| | 0.57 | 0.65 | 0.59 | *NJ* |
| | | 0.72 | 0.67 | *tsinfer* |
| | | | 0.77 | *Relate* |
| | | | | *Singer* |
| **Tv- Variety Topology** | | | | |
| *NJ* | *tsinfer* | *Relate* | *Singer* | |
| | 0.57 | 0.65 | 0.59 | *NJ* |
| | | 0.66 | 0.61 | *tsinfer* |
| | | | 0.73 | *Relate* |
| | | | | *Singer* |
| **Ta - Alternate Topology** | | | | |
| *NJ* | *tsinfer* | *Relate* | *Singer* | |
| | 0.45 | 0.56 | 0.50 | *NJ* |
| | | 0.62 | 0.57 | *tsinfer* |
| | | | 0.71 | *Relate* |
| | | | | *Singer* |
